# Collagenesis Orchestrates Mechanoadaptive Homeostasis of Intervertebral Disc

**DOI:** 10.1101/2023.07.24.550303

**Authors:** Jian He, Sha Huang, Yangyang Li, Yingbo Wang, Pulin Yan, Ou Hu, Peng Lin, Huaijian Jin, Jun Zhu, Liang Zhang, Yaoyao Liu, Qin Qin, Yu Guo, Xiuhui Zheng, Yangli Xie, Lin Chen, Yu Lan, Bing Liu, Peng Liu, Yibo Gan

## Abstract

Intervertebral discs are crucial to spine flexibility and stability during locomotion, but the mechanism of their mechanoadaptation to stress remains inadequately elucidated. We discovered spontaneous collagenesis in discs at the tail base in mice occurring from 14 days post-natal development. These discs termed as type C, differ from typical discs by exhibiting intensive collagen II and reduced aggrecan deposition contributing to the stiffening of the extracellular matrix (ECM), without typical degenerative markers. The reinforced mechanical properties enable type C discs to withstand high mechanical stress during flexion. Further analysis revealed that type C discs experience a turnover in cell composition, with the emergence of Procr^+^ progenitor cells and depletion of notochord cells. We demonstrated the essential role of mechanical stress in type C disc formation, suggesting a mechanoadaptive process rather than a pathological condition. Mechanistically, we identified TRPV4, as pivotal factors in collagen II synthesis in type C discs, highlighting the role of mechanotransduction in this adaptation. Our findings introduce a novel mechanoadaptive process of intervertebral disc by orchestrating collagenesis, advancing understanding of diverse disc functions in spine development and homeostasis, thereby providing insights for leveraging this mechanism to address spinal disorders.

## Introduction

The spine is the central skeletal scaffolding for the body, providing support for locomotion and protecting the spinal cord and nerves. The intervertebral disc (IVD) is the critical unit responsible for bearing mechanical stress to maintain spinal integrity. It is commonly believed that mechanical overload can lead to IVD degeneration (1, 2). Cross-sectional population studies show that degenerative changes in IVD can occur in various percentages of individuals, ranging from 12% in those under 18 years old to 70% in those under 50 years old (3–6). However, more than 95% of these individuals are either asymptomatic or have received a diagnosis for disease condition (7, 8). These discrepancies indicate that the IVD is able to withstand the physical demands of daily activities in most cases, prompting further investigation into the mechanisms that allow the spine to adapt to mechanical stress.

The mouse spine disc shows substantial similarities in aspects of geometry and mechanical properties to those of humans, ensuring its potential in studying the mechanism of human disc adapted to mechanical stress (9, 10). The mouse tail is experiencing sudden tilt perturbations, serving as “dynamic rudder” in body balance (11). Despite the significant and continuous mechanical stress from these demands, it is commonly believed that the mouse disc is able to maintain its homeostasis until old age (12, 13). This raises a scientific question about the fundamental cellular and molecular basis underlying this mechanical adaptation.

Here, we revealed a heretofore unrecognized type of disc featured by spontaneous collagenesis (termed as type C) by constructing spatiotemporal profiles of the mouse spine across the postnatal development. In contrast with the typical one fulfilled with proteoglycan and notochord cells (NCs), type C discs are located at the base of mouse coccygeal spine featured by reduced aggrecan deposition but intensive collagen II content, and a lack of typical degenerative markers. The consequent ECM stiffening leads to reinforced mechanical properties. Single-cell RNA-sequencing analysis revealed that type C discs enriched Procr^+^ progenitor cells as we identified in previous studies (14, 15), showing NC-independent developmental origins. The examination on coccygeal spine after unloading demonstrated the essential role of mechanical stress in formation of type C discs. We further characterized the mechanosensory genes, particularly TRPV4, that regulate the collagen II synthesis in type C discs. These findings provide critical insights into the mechanoadaptive mechanisms of IVDs that are essential for preserving disc homeostasis, which are crucial for devising strategies to improve spinal function and potentially alleviate relevant disorders.

## Results

### Spatial-temporal profiling of the mouse spine delineates the unparalleled development of coccygeal discs in the tail base

To investigate the impact of mechanical stress on the postnatal development of the spine, Micro-CT imaging was utilized to examine the structural differences in vertebrae of one-month-old mice (Supplementary Figure 1A). The results indicated that vertebrae ranging from the second coccygeal vertebra (Co2) to Co8 exhibited an hourglass shape, increased bone volume and moment of inertia, suggesting their enhanced strength in resisting mechanical loads (Supplementary Figure 1, B and C)(16, 17). Furthermore, we found a reduced relative disc height index and disc convexity index in these segments (Supplementary Figure 1D). Considering the involvement of coccygeal segments in tail movements in mice, it is reasonable to hypothesize that this spine segment experiences intense mechanical stress(18, 19).

We then profiled the histological characteristics of spine segments spanning from Co2 to Co8 during postnatal development (Figure 1A). We identified substantial variations in disc morphology at postnatal day 7 (P7) and beyond (Figure 1B). Notably, Co4/5 and Co5/6 showed lower disc heights after P28 (Figure 1C and Supplementary Figure 1E). We observed significantly higher histological scores in Co4/5 and Co5/6 discs from P28 to 12 months (Figure 1D), primarily contributed by changes in features of the NP and annulus fibrosus (AF) (Supplementary Figure 1F)(20). Subsequently, Thompson grading (21, 22) revealed that a higher proportion of NP and AF in Co3/4, Co4/5 and Co5/6 were categorized as grade 3 during postnatal development, indicating a dramatic structural change in these discs (Figure 1E). We then used T2-weighted magnetic resonance imaging (MRI) to examine the radiological features. The results showed notable reductions in MRI index and T2 relaxation time at the Co3/4, Co4/5 and Co5/6 levels compared to Co2/3, Co6/7, and Co7/8, indicating decreased hydration in these discs (Figure 1F and Supplementary Figure 1G). To sum up, the comprehensive histological profile of the mouse spine revealed distinct morphological features in Co3/4, Co4/5, and Co5/6, displaying lower disc heights and higher histological scores, suggesting a unique developmental program for these discs.

**Figure 1.**
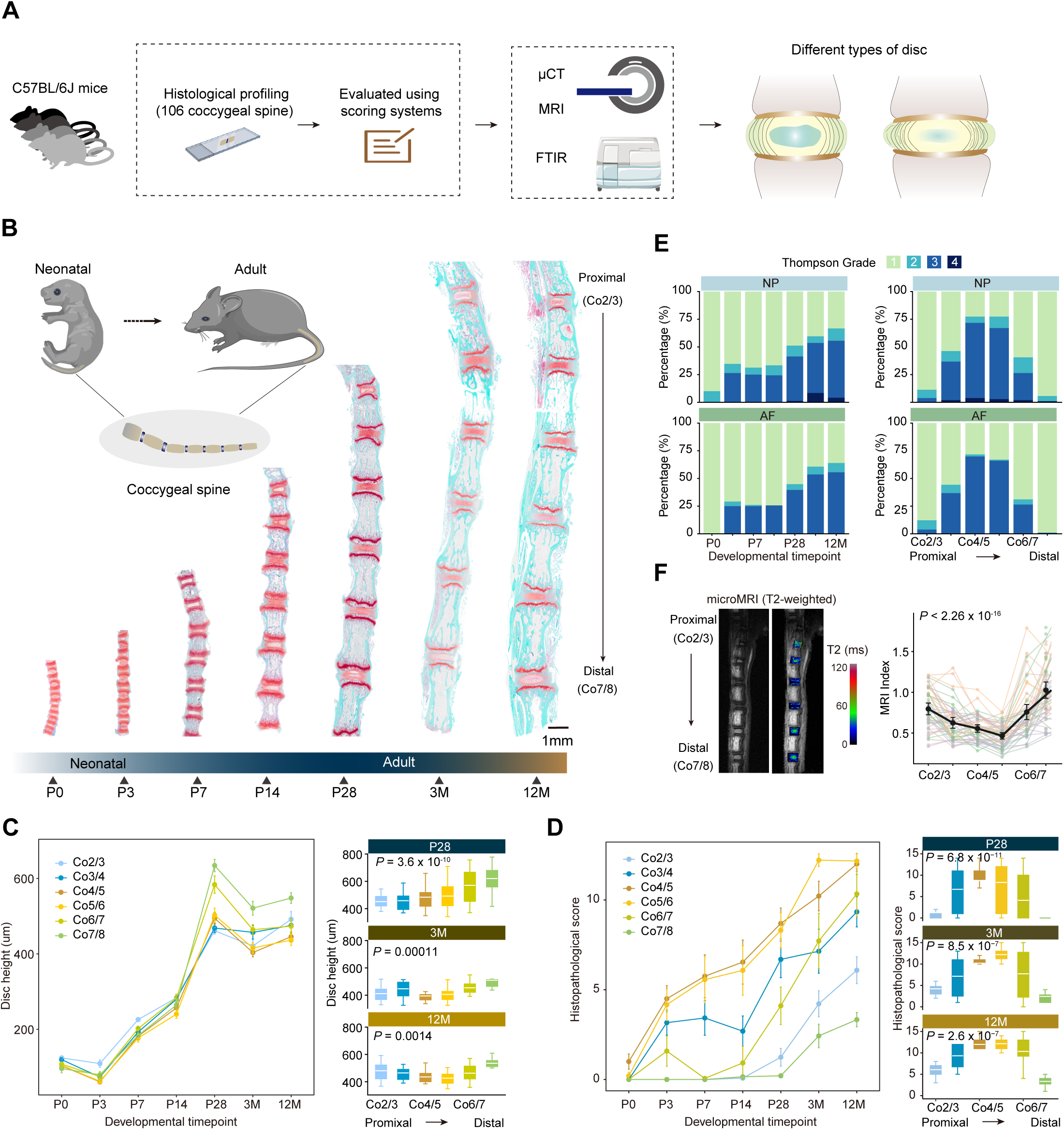
Histological profiling of mouse coccygeal spine identifies spatiotemporal characteristics of the disc. (**A**) Schematic illustrating the design of the profiling study. A total of 106 coccygeal spines from neonatal to mid-aged C57BL/6J mice were collected for histological profiling and evaluation using a scoring system. The integration of micro-CT (μCT), magnetic resonance imaging (MRI) and Fourier transform infrared spectrometer (FTIR) facilitates the identification of distinguishing characteristics of two types of discs. (**B**) Representative sagittal views of safranin O/Fast green (SOFG) staining on coccygeal spine spanning from Co2 (proximal) to Co8 (distal) across various postnatal developmental stages. The samples include P0 (postnatal day 0, N = 10), P3 (N = 12), P7 (N = 16), P14 (N = 13), P28 (N = 29), 3M (3-month-old, N = 14), and 12M (N = 12). Cartilage is depicted in red, while fibers and bones are shown in blue. The longitudinal growth of the spine is accompanied by fibrous and osteogenic deposition in vertebrae and chondrogenic deposition in discs. (**C**) The heights of discs from Co2/3 to Co7/8 levels were measured at various developmental time points. Data are mean ± S.E.; Right, detailed comparison of disc levels at P28, 3M and 12M. The central line represents the median, boxes indicate 25^th^–75^th^ interquartile range and whiskers 1.5*interquartile range. *P* values are determined by Kruskal-Wallis test. (**D**) Histopathological scores of discs from Co2/3 to Co7/8 levels were calculated at various developmental time points. Data are mean±S.E.; Right, detailed comparison of different disc levels at P28, 3M and 12M. *P* values are determined by Kruskal-Wallis test. (**E**) Grade distributions of nucleus pulposus (NP, top) and annulus fibrosus (AF, bottom) in discs at various developmental time points (left) and anatomic levels (right), using a modified Thompson grading system reliant on the SOFG staining. (**F**) Representative sagittal views (left) of a 7.0T microMRI image depict a T2-weighted image and relaxation time of the mouse coccygeal spine. Discs from Co2 to Co8 are labeled with gray values and a pseudo-color map indicating the water content in the tissues. MRI index (right) values of mouse coccygeal discs from Co2/3 to Co7/8. *P* value is determined by Kruskal-Wallis test. Segmented line in matching colors denotes discs from the same mouse (N = 47). The dark point represents the means and the line indicates 95% confidence interval.

### Characterization of collagen-abundant discs in the mouse tail spine

Next, we analyzed the macroscopic characteristics of Co3/4, Co4/5 and Co5/6 discs. Strikingly, these discs exhibited a non-gelatinous NP and indistinct borderlines between the NP and AF (Figure 2A). In contrast, the Co2/3, Co6/7 and Co7/8 discs demonstrated a typical appearance characterized by a well-defined NP filled with water content (Figure 2A). Morphological changes were evident in the Co4/5 disc, as indicated by the loss of NP/AF demarcation, reduced NP cell count, and presence of clefts (Figure 2B). Polarized light imaging revealed a higher deposition of collagen fiber in the NP of Co4/5 disc, which was absent in the NP of Co7/8 disc (Figure 2, C and D). Spatiotemporally, this collagen accumulation occurred in Co4/5 and Co5/6 discs since postnatal day 14 (Supplementary Figure 2, A and B). Quantitative analysis demonstrated a similar composition of collagen fibers in the AF of both discs (Figure 2E and Supplementary Figure 2C), suggesting that structural alterations and collagen fiber accumulation primarily occurred in NP.

**Figure 2.**
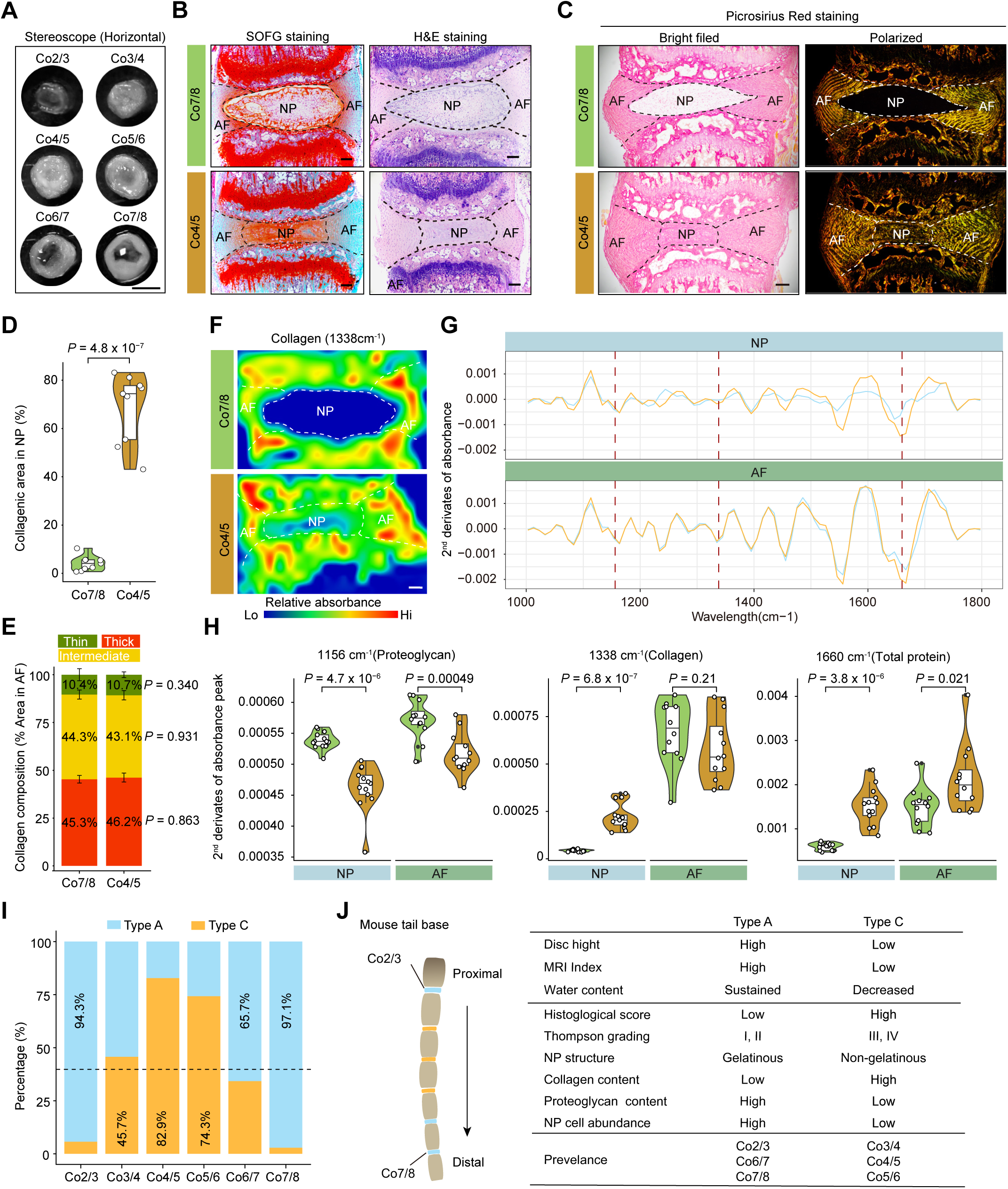
Multilayer-characterization of collagen-abundant disc in mouse tail base. (**A**) Representative horizontal sections of discs ranging from Co2/3 to Co7/8, in which NP region of Co2/3, Co6/7, and Co7/8 exhibit gelatinous and enriched water content, while those of Co3/4, Co4/5, and Co5/6 exhibit non-gelatinous and decreased water content. Scale bar, 1 mm. (**B**) Representative sagittal section of SOFG and hematoxylin-eosin (H&E) staining showed the distinct morphology of Co7/8 and Co4/5 discs. Dashed lines indicate the boundary between NP and AF. Scale bar, 100 μm. (**C**) Representative picture of Picrosirius Red staining on Co7/8 and Co4/5 discs under bright field (left) and polarized light (right). The collagen fiber thickness is denoted by green (Thin), yellow (Intermediate), and red (Thick). (**D**) Violin plots display the collagenic area proportions identified by Picrosirius Red staining in NP from two group sets of discs (N = 10 for each group set). Co4/5 disc with NP fulfilled with collagen fiber. *P* values of significance are determined by two-tailed Student’s t-test. Each point represents the average value from discs in the same group set within individual mice (N = 10). (**E**) Stacked bar charts show the collagen fiber composition in AF of Co7/8 and Co4/5 discs. *P* values of significances between two types of discs are determined by two-tailed Fisher’s exact tests. (**F**) Representative FTIR images of Co7/8 and Co4/5 discs. The color gradient indicates the relative absorbance levels of collagen (1338 cm^-1^). (**G**) Spectrum diagram of average second derivatives of absorbance peak across wavelengths from 1,000 cm^-1^ to 1,800 cm^-1^ in the NP region (top) and AF region (bottom) from Co7/8 and Co4/5 discs. Derivatives at 1156 cm^-1^ (proteoglycan), 1338 cm^-1^ (collagen) and 1668 cm^-1^ (total protein) are indicated by dashed lines. (**H**) Violin plots show the derivatives at three wavelengths of NP or AF from two type of discs (N = 10 for each group). *P* values of significance are determined by two-tailed Student’s t-test. Points represent values at 3 detection points from an individual disc (N = 5). (**I**) Stacked bar charts show the frequency of two types of discs across Co2/3 to Co7/8. The types of discs are identified according to the SOFG staining for each sample, resulting in type A (aggrecan-abundant) and type C (collagen-abundant) discs. The dashed line indicates the average proportion of type C discs in all samples (N = 35). (**J**) Summarized table comparing distinct characteristics in type A and type C discs.

To determine the differences in ECM composition, we used FTIR spectroscopy to assess the protein context in Co7/8 and Co4/5 discs. The results revealed significantly higher collagen synthesis in NP and AF of Co4/5 disc (Figure 2F). Concurrently, we observed proteoglycan degradation alongside an increase in collagen and total protein content in NP of Co4/5 disc (Figure 2, G and H). A minor elevation in collagen content was noted in the endplate (EP) of Co4/5 disc (Supplementary Figure 2, D and E). The differences in NP and AF could be detected as early as P14 (Supplementary Figure 2F). Our findings confirmed the prevalence of collagen deposition in the Co3/4, Co4/5 and Co5/6 discs. These distinct variances in the ECM led to the hypothesis of a novel disc type in the coccygeal spine, designated as collagenic disc (type C), in contrast to the typical aggrecan-enriched disc morphology (type A). The prevalence rates of type C discs range from 45.7% to 82.9% at Co3/4, Co4/5, and Co5/6 levels (Figure 2I). Therefore, our findings illustrate the presence of two distinct types of discs in the mouse tail, each exhibiting unique characteristics that imply diverse functions within the spine (Figure 2J).

### Type C disc presents an ECM remodeling instead of degenerative consequences

We then conducted RNA-seq to compare the transcriptomic differences between type A and type C discs. PCA clearly distinguished the two types of discs, revealing 255 significantly differentially expressed genes (DEGs) for type C and 424 DEGs for type A discs (Figure 3, A and B, and Supplementary Table 1). Strikingly, the core matrisome term with collagens was found to be significantly enriched in the signature genes of type C discs (Figure 3C). Specifically, *Col2a1* but not *Col1a1*, is highly expressed in type C discs, while *Acan*, encoding a proteoglycan, is highly expressed in type A discs (Figure 3D). Immunostaining analysis confirmed the collagen II deposition and aggrecan degradation particularly in the NP of type C discs (Figure 3E). Additionally, signature genes of NCs such as *T*, *Krt8*, *Krt18*, and *CD24a* (23, 24), were significantly downregulated in type C discs compared to type A discs (Figure 3F). Functional enrichment analysis revealed that the upregulated DEGs in type A discs were associated with cell adhesion and notochord morphogenesis, whereas those in type C discs were linked to cellular calcium ion homeostasis, and cell chemotaxis (Supplementary Figure 3, A and B). Gene set variation analysis (GSVA) confirmed that type C discs displayed a higher level of chondrocyte differentiation and a lower level of oxidative phosphorylation (Figure 3G). Interestingly, Weighted Gene Co-Expression Network Analysis (WGCNA) identified a module correlated with mechanosensation (*Piezo1* and *Trpv4*) in type C discs (Supplementary Figure 3, C to E).

**Figure 3.**
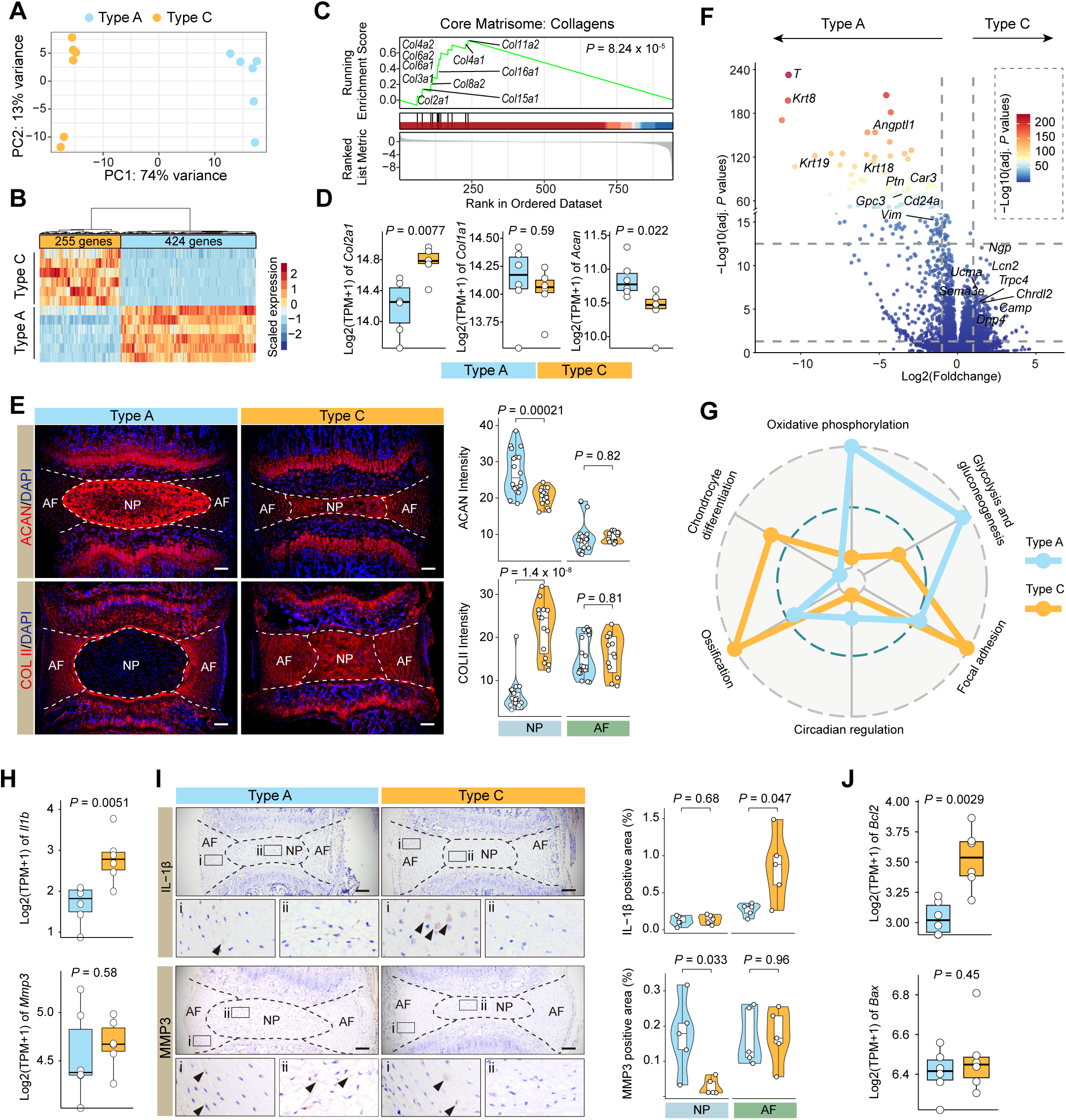
Type C disc does not feature the degeneration characteristics. (**A**) PCA of type A and type C discs (N = 6 for each group). The PC1 (explaining 74% of the variance) and PC2 (explaining 13% of the variance) are set as the axes. (**B**) Heatmap shows the expression (column scaled) of significant DEGs between the two types of discs. DEGs are hierarchically clustered into two branches based on the expression levels. (**C**) GSEA analysis of core matrisome term (Collagens) in upregulated genes of type C discs. Enriched genes in term are labelled. (**D**) Expression levels of *Col2a1, Col1a1* and *Acan*. Box indicates 25^th^, median (middle line), 75^th^ quartiles, and whiskers indicate 5^th^ and 95^th^ percentiles and single points indicate outliers. *P* values of significance are determined by the Wilcoxon rank-sum test. (**E**) Representative immunofluorescent staining of ACAN and COL II in type A and type C discs (left). Quantification of signaling density in NP region and AF region (right). Scale bar, 100 μm. *P* values were calculated with two-tailed Student’s t-tests. (**F**) Volcano map shows the log2-transformed foldchanges and negatively log10-transformed adjusted *P* values for all genes, indicated by the color gradient. Curated significant DEGs are highlighted. (**G**) Radar map shows the performance of six gene sets associated with the indicated functions and metabolic pathways in the two types of discs. (**H**) Expression levels of inflammatory factor gene *Il1b* and degenerative marker gene *Mmp3*. (**I**) Representative immunohistology staining of IL-1β and MMP3 in type A and type C discs. Scale bar, 100 μm. Proteins are brown, and nuclei are blue. *P* values were calculated with Student’s t-tests. (**J**) Expression levels of apoptotic genes *Bcl2* and *Bax*.

To exclude the possibility of early onset of degeneration in type C discs, we conducted a comparative analysis of key markers associated with IDD (25). Our results revealed that levels of IL-1β, a pro-inflammatory cytokine, were expressed at a lower level in NP and elevated in the AF of type C discs (Figure 3, H and I). The expression level of MMP3, a matrix metalloproteinase involved in ECM degradation, was even lower in NP of type C discs (Figure 3, H and I). Notably, typical degenerative characteristics observed in other mouse models were not evident in type C discs (Supplementary Figure 3F) (26–29). Furthermore, IDD-related genes in ECM homeostasis, autophagy, apoptosis and oxidative stress did not exhibit distinct patterns between type A and type C discs (Supplementary Figure 3G). In addition, the heightened expression of *Bcl2* while *Bax* remains unchanged in type C discs indicates an anti-apoptotic response in type C discs (Figure 3J). Collectively, our results highlight ECM remodeling from aggrecan to collagen II in NP of type C discs without manifesting typical degenerative characteristics.

### Type C disc exhibits superior mechanical properties in response to stress

To evaluate the mechanical properties of these two types of discs, we used atomic force microscopy (AFM) to probe the surface characteristics and mechanical response by applying a controlled force (Figure 4A). Our analysis indicated higher viscoelasticity in NP, which was characterized by a tortuous force-displacement curve (Figure 4B). The increased hysteresis during the approaching-retraction cycle reflected enhanced stiffness in type C discs, suggesting a more robust and resilient mechanical structure (Figure 4, B and C).

**Figure 4.**
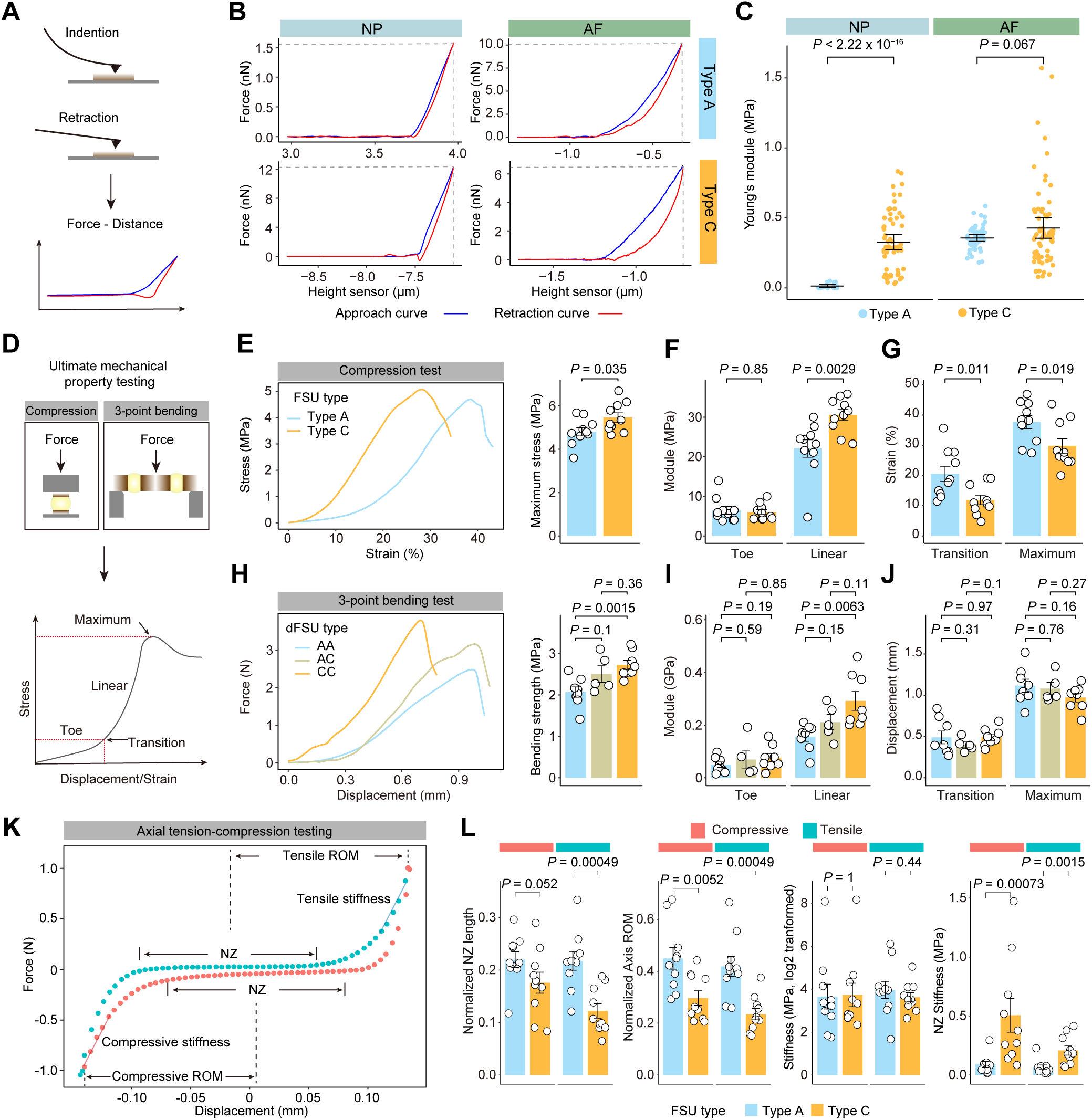
Mechanical properties of type C discs. (**A**) Schematic of AFM analysis on the horizontal surface of discs. Two processes including indentation and retraction are conducted and recorded to generate force-distance curves. (**B**) Representative force-distance curves were detected in the NP region (left) and AF region (right) from type A (top) and type C (bottom) discs. (**C**) The stiffness reflected by Young’s modulus was calculated from each test. Data are shown with mean and 95% confidence interval. Two-tailed Student’s t-tests are used to determine the *P* values. The point represents 25 tests from the individual sample (N = 4). (**D**) Schematic of mechanical property testing. In compression test, FSU consists of the entire disc and adjacent vertebrae. The FSU bears the force from the load cell. In 3-point bending test, dFSU comprises a vertebra sandwiched by two discs. The load cell exerts a force on the middle of the dFSU. The force and displacement are meticulously recorded during the test. (**E**) Representative stress-strain curves detected in compression test on type A and type C discs (left). Quantification of maximum stress in each test (right, N = 10 for each group). (**F**) Quantification of the toe and linear region modulus (N = 10 for each group). (**G**) Quantification of transition and maximum strains. Data are shown as means ± S.E. (N = 10 per group). Significant differences between groups were assessed via two-tailed Student’s t-tests. (**H**) Representative stress-strain curves detected in 3-point bending test on type A and type C discs (left). Quantification of maximum stress in each test (right, N = 10 for each group). (**I**) Quantification of the toe and linear region modulus (N = 10 for each group). (**J**) Quantification of transition and maximum strains. Data are shown as means ± S.E. (N = 10 for each group). Significant differences between groups were assessed with the Wilcoxon rank-sum test. (**K**) Representative of last force-displacement curve. Red dots and blue dots indicate compressive and tensile process respectively. NZ, neutral zone; ROM, range of motion. (**L**) Quantification of NZ length (normalized to FSU length), axis ROM (normalized to FSU length), stiffness, and NZ stiffness in compressive and tensile process. Data are shown as means ± S.E. (N = 10 for each group). Significant differences between groups were assessed with the Wilcoxon rank-sum test.

In order to assess the elastic properties under extreme stress, we conducted compression and three-point bending tests on functional spine units (FSUs) with two types of discs (Figure 4D). The compression test revealed significantly higher maximum stress and linear region modulus in FSUs with type C disc, indicating their greater resistance to elastic deformation compared to type A discs (Figure 4, E and F). Interestingly, the lower strains at both the transition and maximum points in FSUs with type C disc suggested their restricted flexibility (Figure 4G). Three-point bending tests showed that double FSUs (dFSUs) with two type C discs exhibited higher bending strength and linear region modulus compared to those with two type A discs (Figure 4, H to J). To capture mechanical dynamics during reciprocating locomotion, we then conducted cyclic axial tension-compression test (Figure 4K). FSUs with type C discs showed a notably reduced length of neutral zone (NZ) in the tensile curve and axis range of motion in both compressive and tensile curves, aligning with their decreased disc height and limited flexibility as observed in previous analysis (Figure 4L). The higher NZ stiffness in FSUs with type C discs, when compared to that with type A discs, suggests the capacity to withstand higher stress (Figure 4L). Notably, the anticipated rise in compressive stiffness, often associated with disc degeneration(22), was absent in type C discs (Figure 4L). Taken together, these results suggest that type C discs possess enhanced mechanical properties to endure mechanical stress, albeit at the cost of flexibility, providing a physiological basis for adapting to demanding mechanical conditions.

### scRNA-seq analysis reveals the cellular alterations and distinct developmental origin of type C disc

We performed single cell RNA-seq (scRNA-seq) analysis to investigate the cellular heterogeneity of two distinct types of discs (Supplementary Figure 4A). Integrated analysis revealed nine cell clusters based on the expression of canonical marker genes, encompassing NCs (cluster 1), four subsets of chondrocyte (clusters 2-5, RegC, HyperC, FibroC and CycC, representing regulatory, hypertrophic, fibrotic and cycling chondrocytes respectively), osteoprogenitor cells (cluster 6, OP), disc progenitor cells (cluster 7, PC), as well as endothelial and blood cells (clusters 8 and 9) (Figure 5, A and B, Supplementary Figure 4B and Table 2). Cell composition analysis revealed an abundance of NCs expressing Brachyury (encoded by *T*) in type A discs, whereas PCs expressing EPCR (encoded by *Procr*) which have been identified in human and goat discs (14, 15), emerged in type C discs (Figure 5, C and D). Furthermore, deconvolution of bulk transcriptomic dataset confirmed the scarcity of NCs in type C discs (Supplementary Figure 4C). Notably, the residual NC in type C disc exhibited elevated expression of collagen-related genes, such as *Col1a2*, *Col2a1*, and *Col3a1*, which enriched processes related to collagen biosynthesis (Figure 5E and Supplementary Figure 4D). PCs in type C discs showed increased expression of *Dpp4* and *Thy1*, signature genes associated with stem and progenitor cells (Figure 5E) (30, 31).

**Figure 5.**
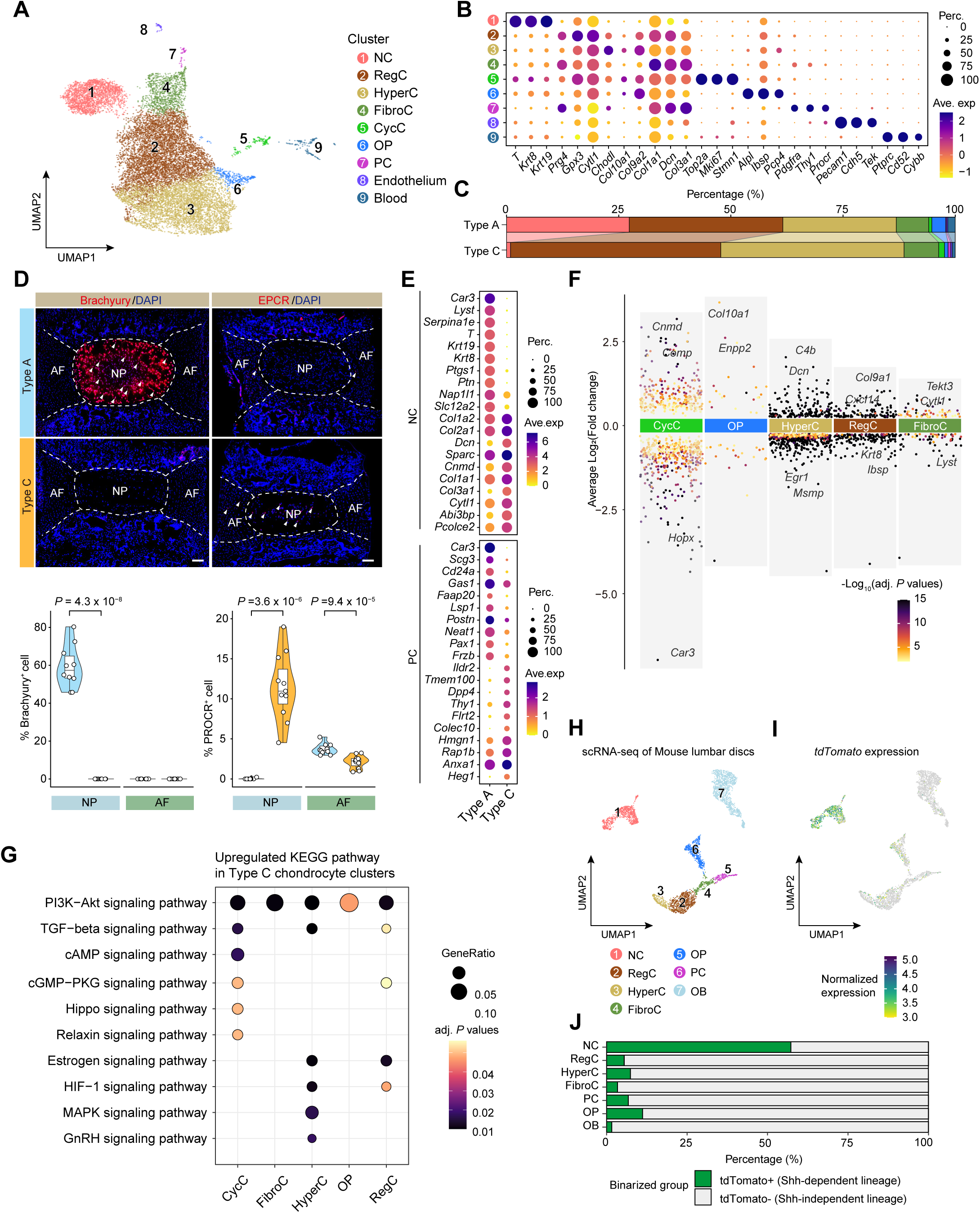
scRNA-seq analysis reveals the cellular alterations and enhanced chondrogenesis in type C discs. (**A**) The Uniform Manifold Approximation and Projection (UMAP) plot depicts the cell clusters identified in two distinct disc types, including Notochord cells (NC), regulatory chondrocytes (RegC), hypertrophic chondrocytes (HyperC), fibrotic chondrocytes (FibroC), cycling chondrocytes (CycC), osteoprogenitors (OP), and progenitor cells (PC). (**B**) Expression of marker genes was used to identify the cell clusters. Color gradient indicates the average gene expression in clusters. Dot size indicates the percentage of gene expression within each cluster. (**C**) Percentages of each cell cluster in type A and type C discs. (**D**) Representative immunofluorescent staining of NC marker Brachyury (*T*) and PC marker EPCR (*Procr*) in type A and type C discs (top). Quantification of T^+^ cell and PROCR^+^ cells in NP and AF region (bottom). Scale bar, 100 μm. *P* values were calculated with two-tailed Student’s t-tests. (**E**) Top 10 differentially expressed genes in NCs and PCs between type A and type C discs. (**F**) Volcano maps of DEGs in chondrocyte clusters between type A and type C discs. Color gradient indicates the P values of each gene. Curated significant DEGs are highlighted. (**G**) KEGG analysis of upregulated genes in type C chondrocyte clusters. The enriched signaling pathway for each cluster are shown in the dot plot. Color gradient indicates the *P* values of enrichment. Dot size indicates the ratio of enriched genes to all genes in the pathway. (**H**) UMAP of disc and VB cell clusters by re-analyzing scRNA-seq dataset of mouse lumbar spine. (**I**) UMAP of tdTomato gene expression in disc and VB cell clusters. (**J**) The expression of tdTomato gene were binarized using QUBIC2 algorithm. The composition of tdTomato^+^ and tdTomato^-^ cells for each cell clusters were calculated.

We then compared the differences in chondrocyte clusters between type A and type C discs. Genes related to chondrocyte differentiation such as *Cnmd*, *C4b*, *Comp*, *Col9a1* and *Cytl1* were found to be significantly upregulated in type C disc (Figure 5F and Supplementary Figure 4, E to G). Moreover, the expression of *Lepr*, *Car1*, and *Adgrg1*, which are specific to chondrocyte progenitors, was significantly higher in type A disc (Supplementary Figure 4H) (32). Pathway analysis revealed enrichment of chondrogenesis-related pathways including PI3K-Akt, TGF-beta and HIF-1 signaling in type C discs (Figure 5G). These results suggest a more pronounced chondrocyte differentiation in type C discs than that in type A discs. Furthermore, we did not find differences in cell senescence or oxidative stress in the chondrocyte clusters, suggesting independence from degenerative and aging processes in type C discs (Supplementary Figure 4I).

To explore the developmental origins of NCs, PCs and chondrocytes, we reanalyzed disc cell clusters from scRNA-seq on Shh-Cre;tdTomato mice, which enable tracking of NCs and their descendants (Supplementary Figure 5, A and B) (33). We used the signature genes of cell clusters in coccygeal disc in our study to identified the cell clusters in lumbar discs (Figure 5H and Supplementary Figure 5, C and D). NCs highly express the *tdTomato* gene, as expected (Figure 5I). Notably, we observed minimal presence of tdTomato^+^ cells in PCs, osteoblasts, and chondrogenic clusters, suggesting their notochord-independent developmental pathways (Figure 5J).

Overall, our scRNA-seq analysis unveiled heightened chondrogenic differentiation for collagen synthesis in type C discs, marked by an increase in PC population and a corresponding reduction in NC population within the NP, showcasing distinct embryonic origins with type A discs.

### Mechanical stress is essential for the formation of type C disc

We postulated that the increased mechanical properties of type C discs are regulated by their elevated mechanical condition. Due to the difficulty in directly measuring mechanical parameters in mouse discs during tail locomotion, we constructed a finite element model (FEM) using Micro-CT images of the lumbo-sacro-coccygeal spine (Figure 6A, Supplementary Figure 6A and Table 3). This method has been widely employed to analyze the mechanical dynamics of tissues under certain conditions (34, 35). By conducting simulations of natural lateroflexion, dorsiflexion and plantarflexion (Supplementary Figure 6B and Video 1 to 3), we identified an escalation in force within the discs ranging from Co2/3 to Co5/6, along with concentrated moments in the Co3/4 to Co5/6 discs (Figure 6, B and C). The assessments on these flexions revealed an elevation in the von Mises stress on the AF and EP of the Co3/4, Co4/5, and Co5/6 discs (Figure 6D and Supplementary Figure 6, C and D). Principal component analysis of stresses categorized discs from different levels into two clusters, including type A (Co2/3, Co6/7, and Co7/8) and type C (Co3/4, Co4/5, and Co5/6) (Figure 6E), consistent with the findings from histological analysis. Notably, NP of type C discs experienced higher mechanical stress compared to type A discs during flexion (Supplementary Figure 6, E and F).

**Figure 6.**
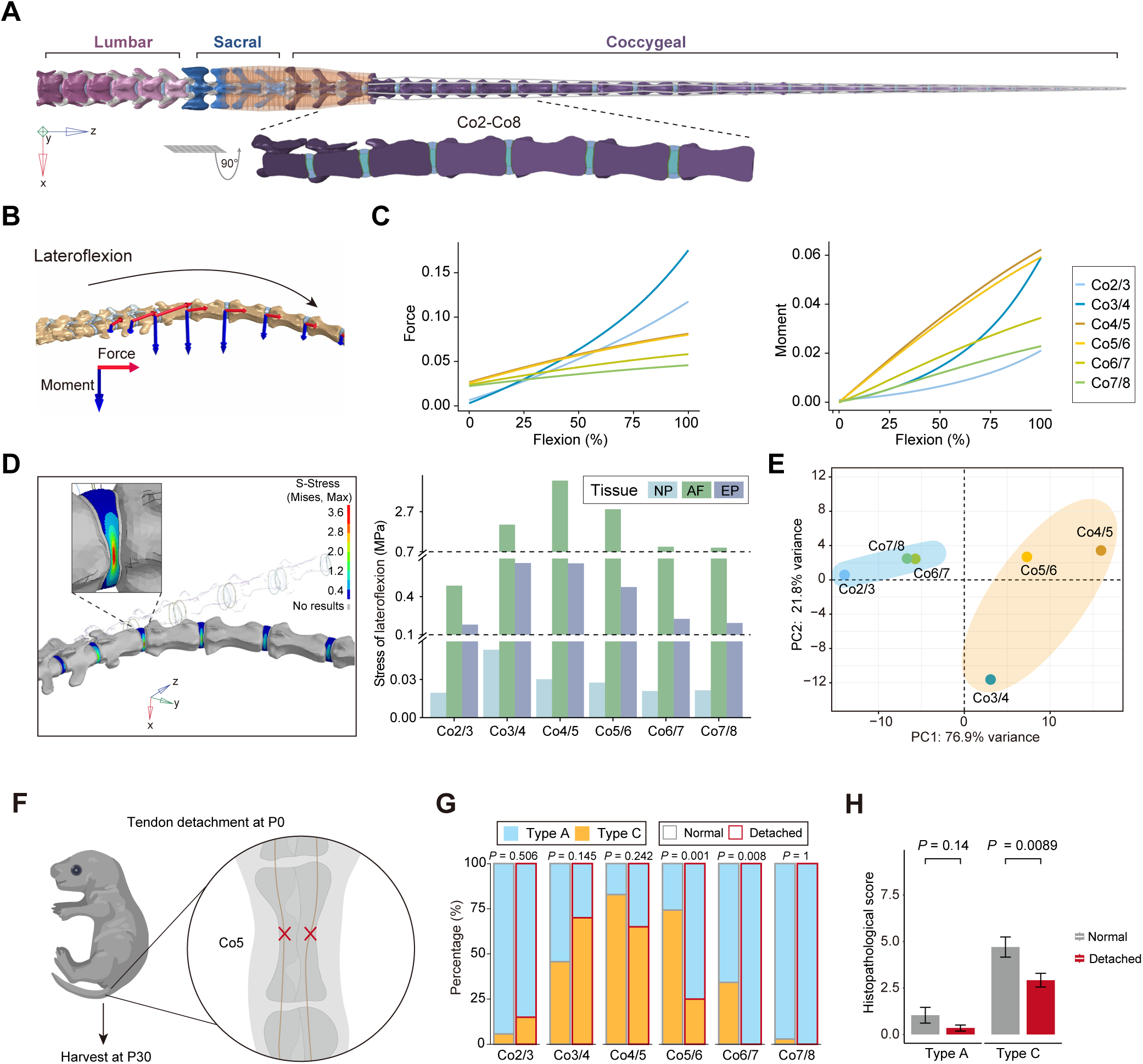
Mechanical stress is essential for the formation of type C discs. (**A**) Vertical view of FEM of mouse vertebrae. Colored vertebral bodies indicate three anatomic parts including lumbar, sacral, and coccygeal vertebrae. The muscle of the tail base is indicated in flesh color and tendons across the caudal vertebra are indicated in gray. A magnified lateral view of Co2-Co8 is shown below. (**B**) Oblique view of FEM of mouse vertebra. The curved arrow indicates the lateroflexion. (**C**) The force (red arrow) and moment (blue arrow) of each disc are captured during the flexion. Dynamics of force and moment of each disc (colored lines) during the lateroflexion. (**D**) View of von Mises stress distributed in AF of mouse tail discs during lateroflexion (left). Magnified view showing the stress distribution of Co4/5 disc. Bar charts show the stress of NP, AF, and EP at the end of lateroflexion. (**E**) Principal component analysis based on stresses of NP, AF, and EP throughout the flexion across Co2/3 to Co7/8 discs. Ellipses showing two group sets of discs. (**F**) Schematic of tendon detachment on mouse tail spine. The tendons surrounding the Co5 were surgically severed at P0 and spine segment specimens were harvested at P30. (**G**) Percentages of two types of discs ranging from Co2/3 to Co7/8 levels. *P* values for the significance of differences in composition between normal control and detached group are determined by two-tailed Fisher’s exact tests (N = 35 for normal mice with sham operation and N = 23 for mice with tendon detachment). (H) Histopathological scores of type A and type C discs from normal control and detachment. Data are mean ± S.E. *P* values are determined by two-tailed Student’s t-tests.

To investigate the necessity of mechanical stress on the formation of type C discs, we performed microdissection to excise the tendon around the tail base at Co5 on the date of birth (Figure 6F). The growth and development of the mouse were not impeded but the tail length was shortened by the tendon detachment, accompanied by decreases in the heights of vertebrae ranging from Co6 to C8 and discs ranging from C5/6 to Co7/8 (Supplementary Figure 7, A and B). On day 30 post-detachment, mice were subjected to treadmill testing to confirm the success of mechanical unloading of the tail (Supplementary Figure 7C and Video 4). Strikingly, we found a significant decrease in the prevalence of type C disc at Co5/6 and Co6/7, which experienced mechanical unloading postnatally (Figure 6G). Accordingly, histopathological scores of type C discs in tendon detachment were markedly lower compared to the normal control with a sham operation (Figure 6H and Supplementary Figure 7D), indicating that mechanical stress is essential for the formation of type C discs.

### TRPV4-mediated mechanoadaptation affects the function of type C disc

The high mechanical stress in type C discs prompted us to inquire about the potential role of mechanical sensing signaling. We observed a significant upregulation of the *Piezo1, Piezo2* and *Trpv4* genes, which are known to be involved in mechanosensitive signaling pathways (36, 37) (Figure 7A and Supplementary Figure 8A). Consistently, a heightened presence of TPRV4^+^ and PIEZO1^+^ cells was predominantly observed in NP of type C discs (Figure 7B), indicating the potential role of mechanosensitive signaling pathways.

**Figure 7.**
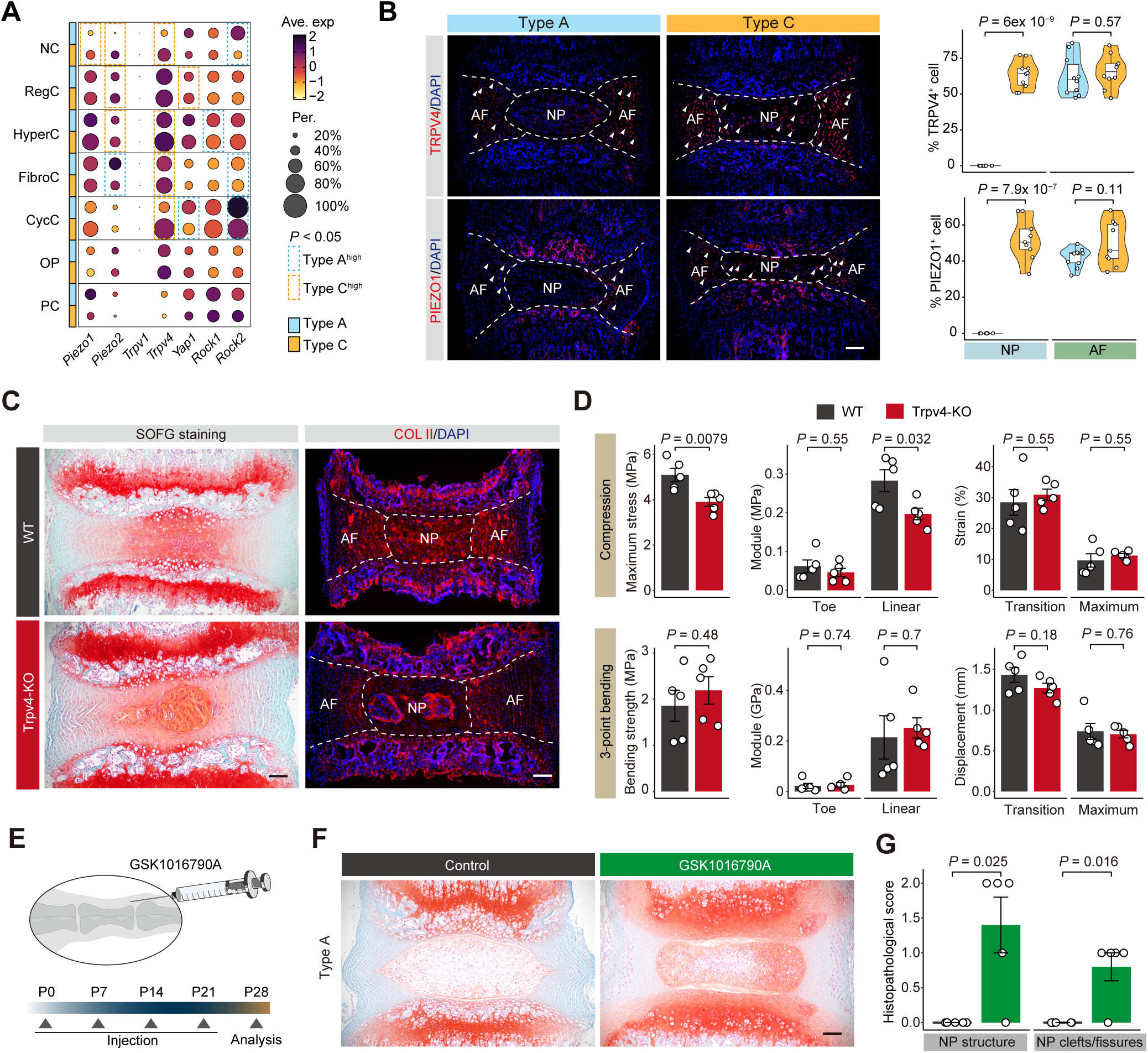
TRPV4-mediated mechanoadaptation affects the function of type C disc. (**A**) Expression of mechano-sensitive genes in each cell cluster. The significance of differences in gene expression between type A and type C discs is determined using two-tailed Student’s t-test. Cell clusters with *P* < 0.05 are boxed in black. (**A**) Representative immunofluorescent staining of PIEZO1 and TRPV4 in type A and type C discs (top). Quantification of positive cells in the NP region and AF region (bottom). Scale bar, 100 μm. *P* values were calculated using Student’s t-tests. (B) Representative SOFG staining and immunofluorescent staining of COL II show the changes in type C disc of Trpv4-ko mice. Dashed lines indicate the border between NP and AF. Scale bar, 100 μm. (**C**) Comparisons of mechanical properties between WT and Trpv4-KO mice assessed by compression test (top) and 3-point bending test (bottom) on type C discs. *P* values for differences between groups were assessed via two-tailed Student’s t-tests (N = 5 for each group). (**D**) Schematic of GSK1016790A (Trpv4 agonist) injection to mouse tail spine. The injection is performed weekly starting from P0 and coccygeal spine segments are harvested at P30. (**E**) Representative SOFG staining shows the morphology changes of NP in type A discs of mice injected with GSK1016790A. Scale bar, 100 μm. (**F**) Histological scores of type A discs from control and injection groups. Data are mean ± S.E. *P* values are determined by two-tailed Student’s t-tests.

To investigate the role of mechanosensitive channel TRPV4 in type C disc, we generated *Trpv4* knockout (*Trpv4*-KO) mice (Supplementary Figure 8B). Compared to their wildtype (WT) littermates, *Trpv4*-KO mice did not exhibit significant differences in general traits, tail length, or disc heights (Supplementary Figure 8, C to E). However, type C discs from *Trpv4*-KO mice showed morphological changes, with obstruction to COL II synthesis in the NP and AF (Figure 7C and Supplementary Figure 8F). Compression and 3-point bending tests revealed a decreased linear modulus response to compression in the FSUs with type C disc in *Trpv4*-KO mice, while their ability to respond to bending remained unaffected (Figure 7D). These findings confirm the crucial role of Trpv4 in ECM remodeling and resulting enhancement of mechanical properties during type C disc formation.

We administered a TRPV4 agonist GSK1016790A to type A discs on postnatal day 0, in an attempt to trigger type C discs during postnatal development. (Figure 7E). Disc treated with the agonist exhibited significantly increased characteristics of type C with collagen deposition, along with increased histopathological scores, compared to the sham control at day 30 post-administration (Figure 7, F and G). Conversely, injections of a PIEZO1 agonist Yoda1 did not induce features of type C disc (Supplementary Figure 8, G and H). These data further emphasize the importance of TRPV4 activation in ECM remodeling and suggest the potential to produce type C discs. In summary, our findings indicate that mechanical stress promotes the formation of type C discs through Trpv4-mediated mechanotransduction.

## Discussion

The intervertebral disc serves as a shock absorber during axial loading, maintaining spinal stability and allowing controlled vertebral movement. By comprehensively profiling the spine in growing mice, we identified a heretofore unidentified disc type (type C) characterized by collagenesis. We consider this to be a mechanoadaptive strategy in response to mechanical stress during spine development. Our findings challenge the traditional view that mechanical stress primarily leads to degeneration or other functional deterioration. This paradigm shift opens up new avenues for deciphering the intricate relationship between mechanoadaptation and its physiological basis, leading to innovative approaches for preventing and treating disc diseases, enhancing spinal stability and overall health.

Mechanical stress resulting from bipedalism is recognized as the primary etiological factor for spinal conditions in humans, notably contributing to the high prevalence of lumbar disc degeneration (1, 2, 38). However, a recent clinical trial study found that engaging in progressive walking can reduce the risk of recurrent low back pain, indicating that the reinforcement by mechanical stress prevents spine dysfunction (39). Our study on mice reveals that type C discs exhibit enhanced mechanical properties to withstand compression and bending stress. The ECM remodeling with increased collagen II and decreased aggrecan within the NP endows the function of type C discs positioned in the highly mechanically stressed area of the mouse tail base. We excluded the possibility of early onset of degeneration as these discs are occurred as early as postnatal day 7 and of lacking typical degenerative characteristics. In contrast, type A discs are common in areas of the spine with lower mechanical stresses, displaying typical characteristics associated with spinal flexibility. This close cooperation is integral for maintaining disc homeostasis while preventing disc from excessive stress (Figure 8). The collagenesis observed in mouse discs may reflect a mechanoadaptive mechanism observed in young human discs (40), which do not result in pathological outcomes under the mechanical loads of daily activities (41). Augmenting collagen deposition, potentially through fibroblast transplantation, could further stabilize the mechanical properties in spine (42), suggesting the mechanical regulation of disc by targeting ECM.

The rodent disc is considered to comprise high retention of NCs that expressing Brachyury (encoded by *T*) in discs throughout their lifespan (9, 43). Nonetheless, our study revealed a distinct disc phenotypes of type C displaying a depletion in NCs. Recent research suggests that the scarcity of T expression is a determining factor in human evolution of tail loss (44), aligning with our concept that the absence of T^+^ NCs in type C discs further represents a mechanoadaptive adaptation in mouse spines. Interestingly, we discovered that type C discs enrich Procr^+^ progenitor cells, a population also identified in healthy human and goat IVDs (14, 15). Our analysis revealed Procr^+^ progenitor cells have a notochord-independent origin, suggesting the distinct developmental paths between type C and type A discs.

To elucidate the underlying mechanism, we established a mouse model with tendon detachment to achieve stress unloading. The reduced frequency and collagenesis in type C discs validate that mechanical stress is essential for the formation of type C discs. Our findings revealed the effectiveness of Pros and Cons of mechanical stress in maintaining disc function, indicating that a delicate balance of forces is crucial for optimal disc health. Given that the mechanical load on the disc varies during mouse tail locomotion (45, 46), further research is needed to quantify the loads associated with different locomotion scenarios, in order to better understand the balance between the “safe window” and “wear and tear” theories (47). Exploring how different mechanical environments impact disc cellular behavior is essential to pinpoint potential therapeutic approaches for enhancing disc resilience and preventing degeneration. This involves studying the molecular pathways activated by mechanical loading and unloading, as well as understanding the role of various cell types in responding to these mechanical changes.

Our transcriptomic analysis suggests that mechanosensation plays a significant role in shaping type C discs. This is evidenced by disruptions in collagen II synthesis observed following knockout of the mechanosensitive gene *Trpv4*. The crucial roles of TRPV4 in regulating IVD mechano-biology have been confirmed (48, 49), with studies showing that inhibiting Trpv4 can prevent IVD degeneration (50–53). Recent research demonstrated that Trpv4 knockout resulted in increased *Acan* mRNA expression, elevated PRG4 protein levels, and reduced histological scores in Co6/7 discs of 4 and 8-month-old mice (54). This effect is consistent with our finding that Trpv4-KO led to the prevention of type C disc characteristics. The roles of TRPV4 in murine discs are diverse, exerting positive effects in developing and healthy discs but exacerbating degeneration in diseased discs(55). This distinction is crucial when devising intervention strategies (56–58). Our study suggests that exogenous activation of TRPV4 by its agonist GSK101 holds promise in promoting the formation of new type C disc, thereby warranting further investigation into its targeting strategy to promote disc homeostasis and prevent spinal disorders.

## Materials and Methods

### Mouse strains

The animal procedures conducted in this study were ethically reviewed and approved by the Animal Ethics Committee of the Third Military Medical University (AMUWEC20223385). C57BL/6J mice were sourced from the Experimental Animal Centers of the Third Military Medical University, SPF (Beijing) Biotechnology Co., Ltd, and Vital River Laboratory Animal Technology Co., Ltd (China). Housing and breeding of the mice took place in a regulated specific pathogen-free (SPF) facility at Daping Hospital, Chongqing, China, under controlled temperature, humidity, and lighting conditions. Trpv4 knockout mice were generated utilizing a CRISPR/Cas9/gRNA targeting strategy to delete exons 3, 4, and 5 of the Trpv4 gene allele (Cyagen, China). Heterozygous parents were mated to generate Trpv4 knockout and wild-type (WT) littermates. The F1 generation was validated by PCR using specific primer pairs: 5’-GGAATGATGGTGATTTTGGCTAAC-3’ and 5’- GGTGGCTATTTAGGGACATGGC-3’ for Trpv4 knockout mice, and 5’- CTACTTTGGTGAGTAGCGGTGGAG-3’ along with 5’-GGTGGCTATTTAGGGACATGGC-3’ for Trpv4 wild-type mice, yielding PCR products that differed in size by 56 base pairs for WT (499 bp) and knockout (555 bp) mice (Supplementary Figure 8B).

### MRI scanning

Forty-seven one-month-old mice were euthanized by receiving an intraperitoneal injection of 0.8% pentobarbital sodium (80 mg.kg^-1^). Mid-sagittal images of all caudal intervertebral discs were qualitatively analyzed using MRI to document the structural changes reflected by water molecules. A 7.0 T micro-MRI machine (BioSpec 70/20 USR, Bruker, Germany) was employed to acquire T2-weighted mid-sagittal images and qualitatively analyze T2 mapping of the discs, using a TurboRARE T2-weighted sequence (repetition time = 4000 ms; echo time = 30 ms; field of view =25×25 mm^2^; slice thickness = 0.4 mm) and T2 mapping sequence (repetition time = 3000 ms; echo times = 9, 18, 27, 36, 45, 54, 63, 72, 81, 90, 99, 108 ms; field of view = 25×25 mm^2^; slice thickness = 0.4 mm). FIJI distribution ImageJ software (v1.54d, National Institutes of Health, USA) was used for manual selection of the region of interest (ROI) for each disc and quantitative analysis of the sagittal T2 image slices. T2 relaxation times were calculated using Image Display and ParaVision (v6.0.1, Bruker, Germany). The MRI index (the average signal intensity) was modified from a previous study to quantify the hydration of the observed disc, with normalization performed against the signal intensity of the upper vertebral bone. The MRI index is defined as:

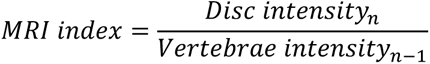

where *n* is the level of the disc. The average signal intensity of the disc or vertebrae was quantified using ImageJ.

### Micro-CT scanning

The specimen of the spine from L1 to the tail end were harvested from one-month-old mice. After removing the skin and soft tissues, we fixed the specimen in 4% PFA. Subsequently, we scanned it using a micro-CT system (Skyscan 1272, Bruker, Germany) with a resolution of 5 μm, 1300 ms exposure time, 65 kV source voltage, and 153 μA source current along with a 0.25 mm aluminum filter. A total of 385 scanning layers were obtained with a 360-degree rotation and 3D reconstructions were generated by the 3-matic Medical software (v17.0, Materialise, Belgium). The width and height for each vertebra were measured along the mid-sagittal plane, and the volume was calculated using the integral algorithm in HyperMesh software (Release 2022, Altair, USA). These scanning layers were then used to establish a finite element model for simulating the motion of the mouse tail. The relative disc height index and disc convexity index were calculated according to a previous study(59). To quantitatively evaluate the disc height, 10 specimens of coccygeal vertebrae from Co2 to Co8 were harvested from one-month-old mice. To maintain the original morphology of the disc, the fixed samples were scanned using a high-resolution micro-CT system (Skyscan 1276, Bruker, Germany) with a pixel size of 10 μm, 1,014 ms exposure time, 50 kV source voltage, and 200 μA source current along with a 0.25 mm aluminum filter. Measurements of volume and moments were performed using CTan software (v1.20). The relative disc height (DH) for each disc was calculated along the mid-sagittal plane according to a previous study(60).

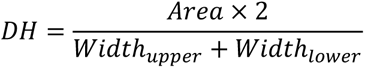

Where the *area* was calculated between the cephalic and endplates and the *width* included the upper and lower edges of disc. Measurements were repeated at least three times to obtain an average for each disc.

### Histological analysis

A total of 106 mouse spines were collected for histological profiling purposes. Specifically, 10 spines at postnatal day 0 (P0), 12 spines at P3, 16 spines at P7, 13 spines at P14, 29 spines at one month (1M), 14 spines at 3M, and 12 spines at 12M were taken. The mice in each group were obtained from at least three littermates. The spines were cut off at the sacral region, and the levels of the coccygeal disc were determined based on the last sacral vertebra. The surrounding connective tissues, including tendon, muscle, and fascia, were carefully removed to obtain the discs and vertebrae. After that, the specimens from Co2-8 were dissected, fixed in 4% paraformaldehyde for 48 hours, then decalcified in EDTA decalcifying solution (R1171, Solarbio) for 20 days before being embedded in paraffin. In the case of disc specimens, tissues were sectioned to the sagittal plane at a thickness of 3 µm in a serial manner, and the gross area of the NP was continuously measured under a microscope. The largest section, or the middle one of the largest sections, was selected for staining analysis and subsequent quantitative assessments. In the case of whole spine specimens, tissues were straightened before fixation to ensure that the middle sagittal views of all discs were aligned at the same focal plane. The sections were stained using hematoxylin and eosin (E607318, Sangon, China), Picro-Sirius Red (G1472, Solarbio, China) for visualization of collagen content or Safranin-O/ Fast Green (G1371, Solarbio, China) for histological assessment. The stained images of the discs were examined under a microscopy system (CX43, Olympus, Japan) and analyzed using FCSnap (V1.1.21486, Olympus, Japan). For the spine from Co2 to Co8, the staining pictures were captured by the VS200 system (Olympus, Japan) and analyzed using OlyVIA software (V3.4.1, Olympus, Japan). The qualification of collagen composition was determined according to a previous study(61). The thickness of fibers in a Picro-Sirius Red staining picture was determined by the value of green, yellow, or red pixels for each point, with green representing thin fibers, yellow representing intermediate fibers, and red representing thick fibers. The region of interest (ROI) was manually selected to distinguish between the NP and AF compartment using ImageJ. To perform Safranin-O/Fast Green staining, the paraffin sections were rehydrated, and then stained with hematoxylin, fast green, and safranin for 5 min successively. Finally, the slides were dehydrated and sealed.

To evaluate morphological changes in coccygeal discs, mid-sagittal sections were scored using two histological reference system. The changes in three components including NP, AF, and EP were determined according to the detailed guidelines by four blinded observers(20, 62). The histological scores for each disc were accumulated from all indicators and averaged by the observers. Additionally, a modified Thompson grading system was applied for evaluating the gross morphological changes in NP and AF(21, 63, 64). Discs were graded into five categories by four blinded observers and the final Thompson grades for each disc were determined by the most voted values. In order to reduce observer variation, intra-observer and inter-observer agreement values were calculated using Fleiss’ Kappa in R function *kappam.fleiss*. Measurements with κ > 0.6 were considered to have substantial agreement and valid for statistical analysis.

### Immunofluorescence staining

For immunostaining, deparaffinized sections were initially treated with Endogenous Peroxidase Blocking Buffer (P0100A, Beyotime, China) for 8 minutes, followed by pepsin retrieval (ZLI9013, ZSGB-BIO, China) for 15 minutes and permeabilization with 1% Triton X-100 (Solarbio, T8200, China) for 15 minutes. Subsequently, the sections were blocked with 5% bovine serum albumin (Sigma, B2064, USA) for 30 minutes. Mouse disc specimens were sectioned at a thickness of 3 µm for immunostaining of Aggrecan (1:100, 13880-1-AP, Proteintech, China), TRPV4 (1:100, PA5-41066, Invitrogen, USA), and PIEZO1 (1:100, 15939-1-AP, ThermoFisher, USA). It is important to note that specimen immunostaining of Brachyury/T (1:50, ab209665, Abcam, UK), EPCR (1:100, ab300523, Abcam, UK), and Col II (1:50, ab185430, Abcam, UK) were embedded in OCT (4583, Sakura) and cryosectioned at a thickness of 10 µm using the Cryostar NX50 (Thermo Scientific, USA). The primary antibody was directly incubated with frozen sections for 6 hours at 4℃ for immunofluorescence staining. Finally, an antifade mounting medium with DAPI solution (Beyotime, P0131, China) was used to visualize the cell nucleus. The immunostaining images were captured using the microscopy system (CX43, Olympus, Japan) and the confocal microscopy system (LSM900, ZEISS, Germany), and analyzed with Zen software (v2.3, ZEISS, Germany) and ImageJ software.

### Immunohistochemistry staining

Mouse disc specimens were immunostained with primary antibodies against IL-1β (1:200, NOVUS, nb600-633, USA) and MMP3 (1:100, A22328, Abclonal, China). The sections underwent antigen retrieval in a sodium citrate solution (65°C for 150 minutes), followed by blocking with an Endogenous Peroxidase Blocking Buffer for 8 min and incubation in a 5% BSA mixture with 0.3% Triton X-100 for 30 minutes. Subsequently, the sections were exposed to the primary antibodies overnight at 4°C, processed using a polymer HRP detection system (PV9001, GB-BIO, China), and visualized with a DAB peroxidase substrate kit (ZLI-9017, ZSGB-BIO, China). After staining with hematoxylin for nuclear visualization, the sections were mounted with neutral gum. ImageJ software was used to quantify the density of the positive signal.

### Bulk RNA sequencing

#### RNA isolation and sequencing

Coccygeal discs from twenty-five one-month-old mice were classified into type A and type C through micro-MRI before dissection. The surrounding ligaments and muscle of the vertebral column were removed to expose the discs. Coccygeal disc tissues (Co2/3 - Co7/8) were collected and placed in separate RNA-free tubes on ice (n = 4 - 6 discs for each library) with RNA later reagent (Invitrogen, AM7021, USA). The total RNA for each library was extracted using the MGIEasy RNA toolkit (v3.0, BGI, China). Quality control and abundance for each sample were evaluated using an Agilent 2100 (USA). Equivalent RNA was reversed to cDNA by RT-PCR and then subjected to amplification. The final adapter-ligated cDNA fragments were sequenced on the MGISEQ-2000 platform in a PE150 manner (BGI, China). Reads were aligned to the mouse reference genome (GRCm38) using STAR software (v2.7.1), resulting in a gene-count matrix. Gene expression was quantified as transcripts per million reads (TPM).

#### RNA-seq data analysis

DESeq2 (v1.34.0) package was employed to analyze RNA-seq data. The data dispersion was estimated using the variance stabilizing transformation (VST) function, and hierarchical clustering was conducted based on the Euclidean distance between samples. Principal Component Analysis (PCA) was used for reduced-dimensional visualization. The differentially expressed genes (DEGs) between the two groups were identified using Wald significance tests with the fitting type set as “mean”. Significant DEGs were filtered based on the criterion of log2Foldchange greater than 1 and an adjusted *P* value less than 0.01. In total, 255 upregulated DEGs and 424 downregulated DEGs in type C discs were retained (table S1).

Gene ontology enrichment was performed on DEGs using enrichGO function in clusterProfiler package (v4.2.2). The enriched terms were clustered and visualized using treeplot function in enrichplot package (v1.18.3). GSVA package (v1.42.0) was used to evaluate the scores of gene sets selected from the Molecular Signature Database (MsigDB v2023.1, https://www.gsea-msigdb.org/gsea/msigdb/). Matrisome characteristics were evaluated based on a previous study(65), and the mean scores for each gene set were visualized using the ggradar package (v0.2).

#### WGCNA analysis

The log2-transformed TPM matrix was utilized for WGCNA analysis (v1.72.1). Genes expressed in at least half of the samples were retained for the automatic network construction and module detection using *blockwiseModules* function, with the power parameter set to 9. The topological overlapped values were calculated for the hierarchical clustering of modules with *minModulesize* set to 50. Modules with dendrogram heights below 0.4 were merged, and correlations of eigengenes for each module were calculated. The memberships of module eigengenes with respect to groups were quantified, and the significance of modules was determined by calculating the *P* values of the memberships.

#### Correlation analysis

The RNA-seq expression matrices (GSE100934, GSE128402 and GSE145649) and microarray matrices (GSE134955) were utilized to compare similarities with RNA-seq data from two types of discs. Prior to the comparison, the datasets were first normalized by transforming the count matrices into TPM matrices based on the gene length. Then, Pearson correlation analysis was performed between samples, and the correlation coefficients were visualized using the pheatmap package (v1.0.12).

### Single cell RNA sequencing

#### Single-cell suspension preparation

Twenty one-month-old mice were first examined under MRI scanning to determine the type of disc in the coccygeal spine ranging from Co2/3 to Co7/8. Then, NP and AF tissues from 30 type A and 30 type C discs were harvested and cut into 1 mm^3^ pieces in DMEM medium with 10% fetal bovine serum (FBS, Gibco) and 1% Penicillin-Streptomycin solution (P/S, SV30010, Hyclone) for preparation of single cell suspension respectively. Briefly, the ground disc tissues were digested in DMEM supplemented with an enzyme mixture including 0.2% (w/v) pronase (P0652, Sigma-Aldrich), 0.2% (w/v) collagenase P (11213865001, Sigma-Aldrich) and 100 U/ml hyaluronidase (H8030, Solarbio). The digestion was incubated at 37℃ for 40 minutes until the tissues were completely dispersed. The digestion was incubated at 37℃ for 2 hours until the tissues were completely dispersed. PBS with 2% FBS was used to terminate the digestion and filtered through 40 μm filters with a volume ratio of 1:1. Subsequently, the cell suspension was treated with 1× Red Blood Cell Lysis Solution (130-094-183, Miltenyi). Cell viability was tested by AO/PI staining using the Rigel S2 Cell Counter (Countstar, China).

#### Library construction, sequencing and preprocessing

scRNA-seq libraries for mouse discs were constructed using a DNBelab C Series Single-Cell Library Prep Set (940-001924-00, MGI, China). The process includes single cell encapsulation, mRNA capture, reverse transcription, and cDNA amplification in droplet emulsion. Libraries for each sample were indexed and sequenced using a MGISEQ-2000 sequencer (MGI, China). Raw reads embedded in fastq files were preprocessed using the pipeline of DNBelab C Series HT single cell analysis software (https://github.com/MGI-tech-bioinformatics/DNBelab_C_Series_HT_scRNA-analysis-software, V1.2). The mouse reference genome GRCm38 and gene annotation (Release M23) downloaded from GENCODE database were used to align the reads and quantify gene expression. Count matrices were generated after filtering out low-quality reads with default parameters. For quality control, we applied scDblFinder (v1.8.0) to filter out putative doublets for each data set, and cells with more than 1,000 expressed genes and mitochondrial gene counts less than 10% of all counts were retained for downstream analysis.

#### Integration, dimensionality reduction, and clustering

The Seurat (v4.3.0) workflow was used for integrating data set, performing dimensionality reduction, and clustering. Data normalization was performed using the *NormalizeData* function with a *scale.factor* of 10,000. Highly variable genes of the merged data set were calculated using the *vst* method and used to obtain the top 50 principal components (PCs). The *bbknn* function implemented in the bbknnR package (v1.0.2) was used to align batches by using the merged nearest neighbours for cells in each batch, with the *neighbors_within_batch* set to 3. The corrected neighbourhood graph of bbknn was then used to perform uniform manifold approximation and projection (UMAP) for dimension reduction. For unsupervised clustering, the “bbknn” graph was used and the *Findcluster* function was performed using the Louvain algorithm with a resolution of 0.3, resulting in seven major clusters. Clusters with obvious heterogeneity were then subclustered, leading to a final of nine cell clusters. The published scRNA-seq data from murine lumbar spine cells traced using Shh-Cre;tdTomato strategy were analyzed. To isolate the cells labeled by the Shh expression, we reconstructed the gene expression matrix by aligning reads to a customized reference genome containing the *tdTomato* gene sequence. The *tdTomato* expression for each cell were binarized using QUBIC2 algorithm(66), resulting in the *in silico* percentages of tdTomato*^+^* cells in each cell clusters. The *FindAllMarkers* function implemented in Seurat was used to calculate DEGs among different clusters. The Wilcoxon test was performed on each gene, and the adjusted P-value for statistical significance was computed. Genes that met the following criteria were considered signature genes: 1) expressed in a minimum of 20% of either of the two tested populations; 2) at least a 0.25-fold difference (log-scale) between the two tested populations; and 3) an adjusted *P* values less than 0.01. The identity for each cell cluster was manually annotated based on the differentially expressed signature genes (table S2), which were canonical for specific cell types or reported in previous studies(14, 15, 31).

#### Gene ontology and pathway enrichment

Gene ontology terms enriched in NCs and progenitor cells in type C discs were calculated using the clusterProfiler package embedded in the *RunEnrichment* function in the SCP package (v0.5.4, https://github.com/zhanghao-njmu/SCP). DEGs between cell clusters in type A and type C discs were first calculated and filtered with adjusted *P* values less than 0.01. Then, enrichment analysis was performed using the “GO_BP” database. The clusters of genes and top enriched terms were visualized using the *EnrichmentPlot* function in the SCP package. To compare the signatures in cell clusters between the two types of discs, genes related to the inflammatory response (WP222, WikPathways), elastic fibre (R-MMU-1566948, Reactome) and collagen formation (R-MMU-1474290, Reactome) were extracted for evaluation of scores of gene sets using the GSVA package.

#### Deconvolution analysis

The Scissor package was used to validate the dominating cell clusters in type A and type C discs, as described above. Cell proportion reconstruction for each bulk RNA-seq sample was inferred using the BayesPrism package (v2.1.2)(67). The scRNA-seq dataset was set as the reference, and genes encoding ribosome, hemoglobin, actin, mitochondrial genes, and genes on chromosome X and Y were excluded from the deconvolution process. The *outlier.cut* threshold was set to 0.01 and the *outlier.fraction* threshold was set to 0.1, resulting in a cell cluster proportion table for each bulk RNA-seq sample.

### Finite element model

FE model of a mouse spine from L1 to the tail end was constructed. Initially, the mouse tail was scanned using a micro-CT system as described above. The image files in DICOM format were converted into 3D model construction using the image processing software Mimics. The properties of tissues within the tail, such as muscle and tendon, were identified according to H&E staining and previous descriptions(68), and then used to generate 3D surface models of each tissue type. These 3D surface models were imported into HyperMesh. A tetrahedral mesh was generated for each tissue type, with element size optimized to ensure model accuracy while minimizing computational cost. The mesh was checked for quality using built-in metrics and any problematic areas were manually refined. Material properties were then assigned to each tissue type based on values reported in the literature (Supplementary Table 3). Finally, to simulate the flexions using an implicit method, the lumbar spine was fully constrained to prevent rigid body motion. A pre-loading equivalent to 20% of the maximum force during flexion to the musculature in the mouse FEM. A load was applied as a displacement boundary condition, forcing a 2 mm displacement of the tendon at the base of tail. Three actions including lateroflexion, dorsiflexion and plantarflexion were simulated. The forces and moments for each disc and the von Mises stress on the disc components during the flexion were recorded. PCA was performed on the stress values of NP, AF and EP for each disc along the lateroflexion.

### Atomic force microscopy

Freshly isolated disc specimens were embedded in OCT and longitudinally cryosectioned at a thickness of 100 µm. Sections were fixed on the slide using waterproof glue (Pattex) and kept at 4℃ to avoid dehydration. AFM measurements were performed on a Dimension Icon Scanasyst with a Nanoscanpe 6 controller (Bruker, Germany). For the indentation test, cantilevers coated with reflective gold (k nom = 1 Nm^-1^) with a symmetric tip (radius = 20 nm, height = 8 μm) were used (Ready, China). Before scanning, the cantilevers were calibrated utilizing the thermal tune method with a range set at 1-100 KHz and a spring constant set at 1.115 Nm^-1^. The operation was performed using Nanoscope software (v9.70). The Lorentzian (Air) model was chosen to fit the data. For each disc sample, at least three ROIs were selected for the NP and AF regions under an optical microscope. Each ROI included 25 detection points with a 45-nm spacing. A total of six samples including three type A and three type C discs were measured. Data analysis was performed using the Hertzian model in Nanoscope Analysis software (v3.00). Baseline correction was executed on the extend and retract curves separately. To obtain the force-distance curves and Young’s modulus, indentation analysis was performed with the contact point algorithm set as “Best Estimate” and the Max and Min force fit boundary set at 90% and 30%, respectively. Measurement results with an R value of more than 0.75 were considered valid and retained for subsequent comparative analysis.

### Imaging Fourier transform infrared (FTIR) spectrometry analysis

Disc specimens were longitudinally cryosectioned at a thickness of 10 µm and used to perform FTIR imaging analysis on a Nicolet iN10 Infrared Microscope (Thermo Scientific, USA). Briefly, the specimens were placed onto infrared transparent BaF_2_ windows. The transmission collection mode was applied using a “Cooled” detector, with a wavenumber range of detection set between 4,000 cm^-1^ and 400 cm^-1^. A 100 µm length was used to scan the section, and spectral data were collected for four samples from each group using OMNIC Picta software (Thermo Scientific, USA). For each sample, data from at least three detection points in the NP, AF and EP were collected for subsequent analysis using OMNIC software (v9.2.86). Specifically, the transmission rate curves were transformed into absorbance curves for baseline correction and smoothing to remove the noisy peaks. Then, the absorbance data were processed using the prospectr R package (v0.2.6), in which Savitzky-Golay smoothing and second derivative for the absorbance values were calculated using the *savitzkyGolay* function (*m* = 2, *p* = 3, *w* = 11 and *delta.wave* = 2). The resulting derivatives from detection points in one sample were averaged to minimize technical errors, and multiplied by a factor of negative one for positive visualization. The absolute absorbance values for proteoglycan (wavelength = 1,156 cm^-1^), collagen (wavelength = 1,338 cm^-1^) and total protein (wavelength = 1,660 cm^-1^) were extracted for comparative analysis according to previous studies(29, 64).

### Mechanical property measurement

Mechanical property measurements were conducted using a Universal Testing Machine (AI-7000-SU1, GOTECH, China) equipped with a load cell of 500 N mechanical capacity at room temperature. In compression testing, functional spine units (FSUs) comprising disc and adjacent vertebrae were freshly isolated with soft tissues removed. The descending speed rate for displacement was set as 0.1 mm/minute, and testing ceased when real-time force dropped to 30% of the maximum force. Force-Displacement curves were generated and used to derive Stress-Strain curves based on the disc area and height. Lowess Smoothing was applied to reduce the noise in each curve. Mechanical properties such as maximum stress, toe and linear region modulus, transition and maximum strains were calculated. Briefly, for each curve, the transition point distinguishing the toe region and linear region was identified using the segmented R package (v2.0.2). The modulus for these regions indicated by their slopes, were calculated using the *coef* function from fitted linear models of strain and stress. The maximum strain was determined by the peak stress. The disc type was determined according to the histological staining of the test samples. A minimum of 10 samples were tested for each disc type.

In 3-point bending test, double function spine units (dFSU) comprising vertebrae and bilateral discs were segmented, and categorized into dFSU with two type A discs (AA), dFSU with two type C discs (CC) and dFSU with one type A and one type C disc (AC). The spine specimens were placed horizontally on two supports with a rounded surface (R2 = 2 mm), ensuring suspension of the FSUs, with a span distance of 8.0 mm between the lower supports. A metal indenter with a rounded surface (R1 = 2 mm) was centrally positioned between the supports and connected to the load cell. It descended at a rate of 0.1 mm/minute to apply load to the center of the vertebral body. The test concluded when the force in real-time dropped to 30% of the maximum force. Force-Displacement curves were generated, and the bending strength (σ_b_) for each test was calculated using the following formula:

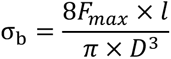

Where the *F_max_* was the maximum force during testing, *l* was the span distance, and *D* was the average transverse diameter of the dFSU. The modulus (E_b_) for the 3-point bending test were calculated using the following formula:

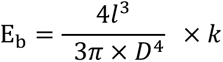

Where the *k* was the slope calculated from the fitted linear models on displacement and force. The toe region and linear region in the curves were determined as described above. At least 5 samples were tested for each category of dFSU.

Cyclic compression and tension tests were carried out on locomotion segments (functional spinal units, FSU) using an Instron 5969 machine with a 100 N load cell capacity at room temperature(69). FSUs were bonded using polymethylmethacrylate-coated fixtures and Loctite 401 adhesive with Loctite SF 770 accelerant, then immersed in physiological saline. Test parameters were set and adjusted based on a previous mouse study(22), with compressive and tensile thresholds at -1 N and 1 N, respectively, a speed of 0.1 mm/s, and an axial pre-load of 0.15 N applied for 1 minute before testing. A total of 10 cycles were performed and the force-displacement curve from the final cycle was fitted using a double sigmoid function, with the 2nd derivative indicating the neutral zone boundaries. The length of the neutral zone, compressive and tensile range of motion, as well as the area of the top and bottom surfaces and height of the FSU, were calculated for statistical analysis. Ten FSUs, including type A or type C discs from 1-month-old mice, were isolated, tested, and stained for disc type identification post-experiment.

### Tendon detachment

The impact of mechanical unloading on the mouse tail disc was assessed through tendon detachment. Briefly, neonatal mouse pups were cryo-anesthetized on a paper-lined ice pack for 5 minutes to induce unconsciousness before the surgical procedure(70). Subsequently, two 5 mm incisions were made in the skin of the tail base along the sagittal direction in both ventral and dorsal aspects to avoid damaging blood vessels. The surrounding tendons were delicately separated and excised using razor blades. It is important to note that the skin was sterilized and did not require suturing due to rapid healing observed the following day. Mice in the sham-operated control group had their tendons left intact. All procedures were conducted under microscopic guidance. The pups were then allowed to recover on an incubator with an infrared heater before being returned to their mother’s cage. To confirm the efficacy of tendon detachment in preventing tail wagging, mice underwent a treadmill test one-month post-surgery (movie S4). Mice that exhibited no tail movement, even when subjected to electrical stimulation, were retained for histological evaluation. Histological scoring was performed on each disc to characterize its type.

### Drug administration

The injection was administered with a 33-gauge needle attached to a Hamilton microsyringe. It was inserted into the skin around the Co6 vertebra and advanced towards the tail base along the sagittal plane. Drugs were manually administered while retracting the needle along the tail axis at a consistent speed. The Trpv4-specific agonist GSK1016790A (HY-19608, MedChemExpress, USA) at a concentration of 10 μM and the Piezo1-specific agonist Yoda1 (HY-18723, MedChemExpress, USA) at a concentration of 500 μM were diluted in 20% SBE-β-CD (HY-17031, MedChemExpress, USA) and 2% DMSO for injection. The dosage of GSK1016790A and Yoda1 for mice of different ages was standardized at 0.1 mg/kg and 1 mg/kg, respectively, according to the recommendations in the instructions. Injections were administered weekly, with a total of four injections performed before harvesting, and subsequent histological staining of the discs ranging from Co2 to Co8.

### Statistical analysis

All statistical analyses were conducted using R (v4.1.3) and software packages compiled in R language. The significance of DEGs in RNA-seq data was assessed using the *result* function in the DESeq2 package with the “Wald” test parameter. Significance was determined using Student’s t-test for the data with homogeneity of variance and normal distribution, and Wilcox-test for the other data. Data points are shown in the barplots and boxplots if the number of replicates is less than 10. Additional statistical analyses are detailed in the corresponding sections of the manuscript. C57BL/6J mice sourced from a minimum of three commercial laboratories were utilized to mitigate the effect of housing environment. Sample sizes were determined to ensure adequate standard deviations of the mean and biologically meaningful differences between experimental groups.

## Data and materials availability

The sequencing data including bulk RNA-seq and scRNA-seq supporting the findings of this study were deposited in the National Omics Data Encyclopedia (https://www.biosino.org/node/index) under the accession number OEP004269 and OEP005392. Mouse sequencing data from published references are available at the GEO with the accession code mentioned in the main text. An interface for scRNA-seq data browsing based on ShinyCell is available online (www.ivdatlas.cn:3838/mIVD) (71). There are no restrictions on data availability. All software and packages including the version are provided in the Method. Custom codes used in the data analysis are available on GitHub (https://github.com/hejian41/mouse_IVD).

## Authors contributions

PLiu, YG, BL and JH conceived and designed the study. YL, LC, and YX provided research consultation. SH, JH, YG, PY, YLi, OH, and P. Lin prepared the animal specimens. YLi. and QQ performed animal breeding. SH, PY, and YW performed immunohistochemistry and immunofluorescence staining. JH and SH conducted the AFM, FTIR, and FSU testing analysis. YG and JH conducted the FEM analysis. YG, SH, and YG performed radiological experiments. JZ, LZ, HJ, YLiu and XZ provided the clinic specimens. JH analyzed and interpreted the sequencing data. JH, YG, and PLiu designed the figures. YG, JH, BL, and PLiu wrote the manuscript. All authors have read and approved the final manuscript.

## Acknowledgments

We thank J. Liu from Jiejun Technique Corporation for his contribution to the FEM construction, W. Ji from Bioengineering College of Chongqing University for assistance during FTIR experiments, and W. Zheng from Olympus Corporation for the help with section staining and imaging. This study was supported by grants from the National Key Research and Development Program of China (2022YFA1103202), the National Natural Science Foundation of China (32270887, 82272507, and 32200654), the Natural Science Foundation of Chongqing (CSTB2023NSCQ-ZDJO008), Postdoctoral Innovative Talent Support Program (BX20220397), the Open Project of State Key Laboratory of Trauma, Burns and Combined Injury (SFLKF202201), Project for Enhancing Innovation of Army Medical University (2023XJS39), and Talent Innovation Training Program at the Army Medical Center (ZXZYTSYS09).

