## Supplementary Figure for "Collagenesis Orchestrates Mechanoadaptive Homeostasis of Intervertebral Disc"

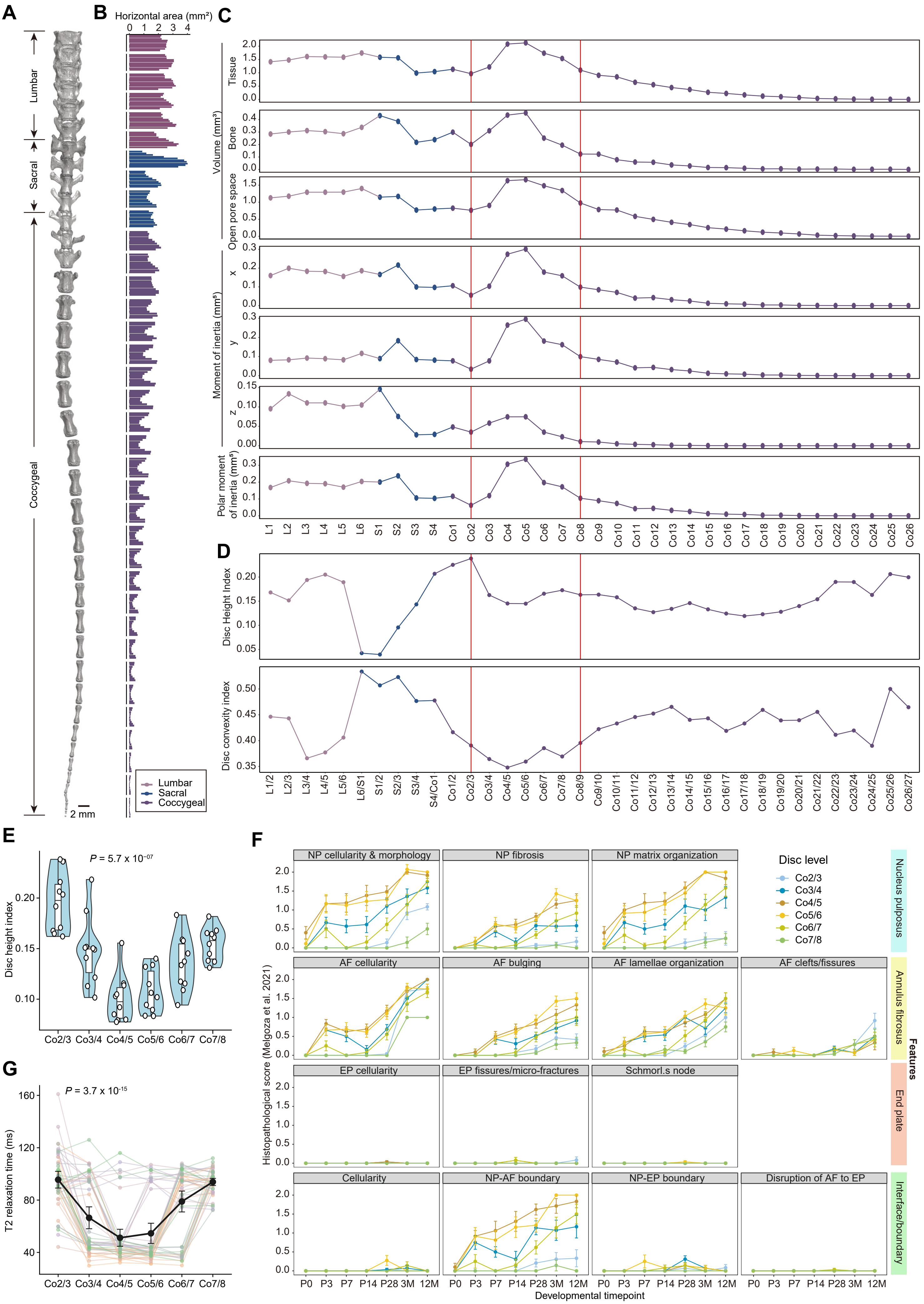

A
Lumbar
Sacral
Coccygeal
2 mm
B
Horizontal area (mm²)
0 1 2 3 4
Lumbar
Sacral
Coccygeal
C
Volume (mm³)
Tissue
Bone
Open pore space
Moment of inertia (mm⁵)
x
y
z
Polar moment of inertia (mm⁵)
L1 L2 L3 L4 L5 L6 S1 S2 S3 S4 Co1 Co2 Co3 Co4 Co5 Co6 Co7 Co8 Co9 Co10 Co11 Co12 Co13 Co14 Co15 Co16 Co17 Co18 Co19 Co20 Co21 Co22 Co23 Co24 Co25 Co26
D
Disc Height Index
Disc convexity index
L1/2 L2/3 L3/4 L4/5 L5/6 L6/S1 S1/2 S2/3 S3/4 S4/Co1 Co1/2 Co2/3 Co3/4 Co4/5 Co5/6 Co6/7 Co7/8 Co8/9 Co9/10 Co10/11 Co11/12 Co12/13 Co13/14 Co14/15 Co15/16 Co16/17 Co17/18 Co18/19 Co19/20 Co20/21 Co21/22 Co22/23 Co23/24 Co24/25 Co25/26 Co26/27
E
P = 5.7 x 10⁻⁰⁷
Disc height index
Co2/3 Co3/4 Co4/5 Co5/6 Co6/7 Co7/8
F
NP cellularity & morphology
NP fibrosis
NP matrix organization
Disc level
Co2/3
Co3/4
Co4/5
Co5/6
Co6/7
Co7/8
Nucleus pulposus
AF cellularity
AF bulging
AF lamellae organization
AF clefts/fissures
Annulus fibrosus
EP cellularity
EP fissures/micro-fractures
Schmorl.s node
End plate
Cellularity
NP-AF boundary
NP-EP boundary
Disruption of AF to EP
Interface/boundary
Features
Histopathological score (Melgoza et al. 2021)
P0 P3 P7 P14 P28 3M 12M
Developmental timepoint
G
P = 3.7 x 10⁻¹⁵
T2 relaxation time (ms)
Co2/3 Co3/4 Co4/5 Co5/6 Co6/7 Co7/8

**Supplementary Figure 1**. **The morphological characteristics of mouse spine identified by microCT and histological assessment.**

(**A**) Reconstructed 3D micro-CT images of vertebral bodies across the mouse spine, encompassing 6 lumbar vertebrae, 4 sacral vertebrae, and 29 coccygeal vertebrae. Scale bar, 2mm.

(**B**) Measurements of vertebral morphologies from micro-CT. The horizontal areas across the spine are measured in 10 cross sections for each vertebra. An abrupt decrease in area is evident in S2, while hourglass dimensions are noted in the vertebrae below Co4.

(**C**) Wave peaks are observed in tissue and bone volumes, moments of inertia in x-y-z dimensions, and polar moment of inertia in the vertebrae spanning from Co2 to Co8, indicating the changes in vertebral structure in response to the mechanical environment.

(**D**) Measurements of disc morphologies from micro-CT. A peak and trough are observed in disc height index and disc convexity index in Co2/3 and Co4/5 discs respectively. This pattern signifies a decrease in disc integrity starting from Co2/3 and a morphological abnormality in Co4/5.

(**E**) Violin plots show disc height index measured from CT image. The central line represents the median, inner boxes indicate $25^{th}$-$75^{th}$ interquartile range and whiskers 1.5*interquartile range. *P* value is determined by Kruskal Wallis test. Each point represents the discs from Co2/3 to Co7/8 in individual mice (N = 10).

(**F**) Four categories of features utilized to evaluate histopathological scores of discs spanning from Co2/3 to Co7/8 levels at various developmental timepoints. Data are mean ± S.E.

(**F**) T2 relaxation time of mouse coccygeal discs from Co2/3 to Co7/8. *P* value is determined by Kruskal-Wallis test. Segmented line in matching colors denotes discs from the same mouse (N = 47). The dark point represents the means and the line indicates 95% confidence interval.

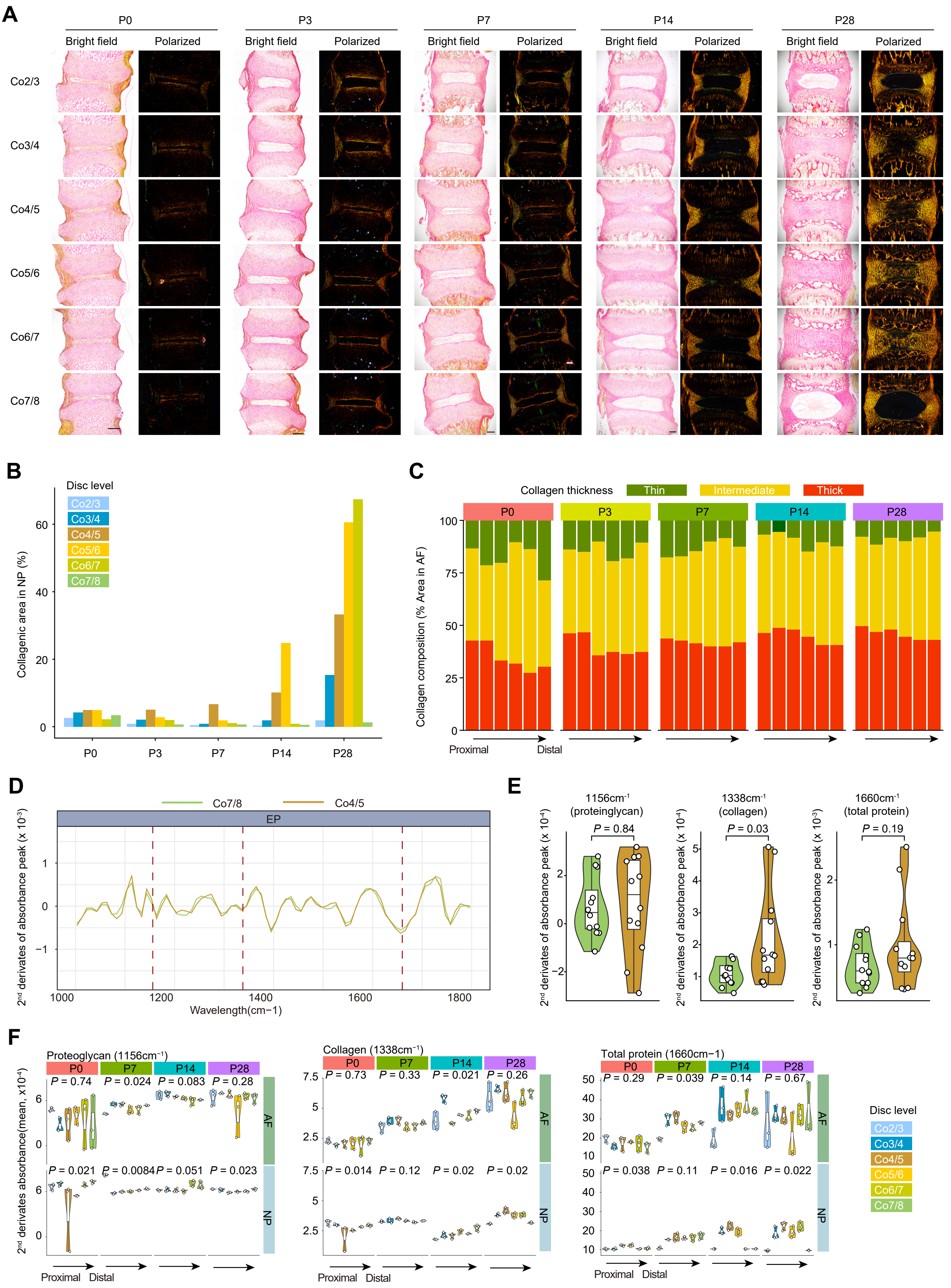

A
P0
P3
P7
P14
P28
Bright field
Polarized
Co2/3
Co3/4
Co4/5
Co5/6
Co6/7
Co7/8
B
Disc level
Co2/3
Co3/4
Co4/5
Co5/6
Co6/7
Co7/8
Collagenic area in NP (%)
C
Collagen thickness
Thin
Intermediate
Thick
Collagen composition (% Area in AF)
Proximal
Distal
D
Co7/8
Co4/5
EP
2nd derivates of absorbance peak (x 10-3)
Wavelength(cm−1)
E
1156cm-1 (proteinglycan)
1338cm-1 (collagen)
1660cm-1 (total protein)
P = 0.84
P = 0.03
P = 0.19
2nd derivates of absorbance peak (x 10-4)
2nd derivates of absorbance peak (x 10-3)
F
Proteoglycan (1156cm-1)
Collagen (1338cm-1)
Total protein (1660cm−1)
2nd derivates absorbance(mean, x10-4)
AF
NP
Disc level
Proximal
Distal

**Supplementary Figure 2. Characterization of collagen context in type C disc at spatiotemporal level.**

(**A**) Picrosirius Red staining on discs from Co2/3 to Co7/8 level under bright field (left) and polarized light (right). The collagen fiber thickness is denoted by green (Thin), yellow (Intermediate), and red (Thick). Scale bar, 100 μm.

(**B**) Collagenic area proportions identified by Picrosirius Red staining in NP from two group sets of discs across postnatal developmental days.

(**C**) collagen fiber composition in AF of discs from Co2/3 to Co7/8 level at various developmental time points.

(**D**) Spectrum diagram of average second derivatives of absorbance peak across wavelengths from 1,000 $cm^{-1}$ to 1,800 $cm^{-1}$ in the endplate (EP) region from Co7/8 and Co4/5 discs. Derivatives at 1156 $cm^{-1}$ (proteoglycan), 1338 $cm^{-1}$ (collagen) and 1668 $cm^{-1}$ (total protein) are indicated by dashed lines.

(**E**) Violin plots show the derivatives at three wavelengths of EP of discs from Co4/3 to Co7/8 discs across postnatal developmental stages. P values of significance are determined by Kruskal-Wallis test (N = 3).

(**F**) Violin plots show the derivatives at three wavelengths of NP and AF of discs from Co2/3 to Co7/8 discs across postnatal developmental days. *P* values of significance are determined by Kruskal-Wallis test (N = 3).

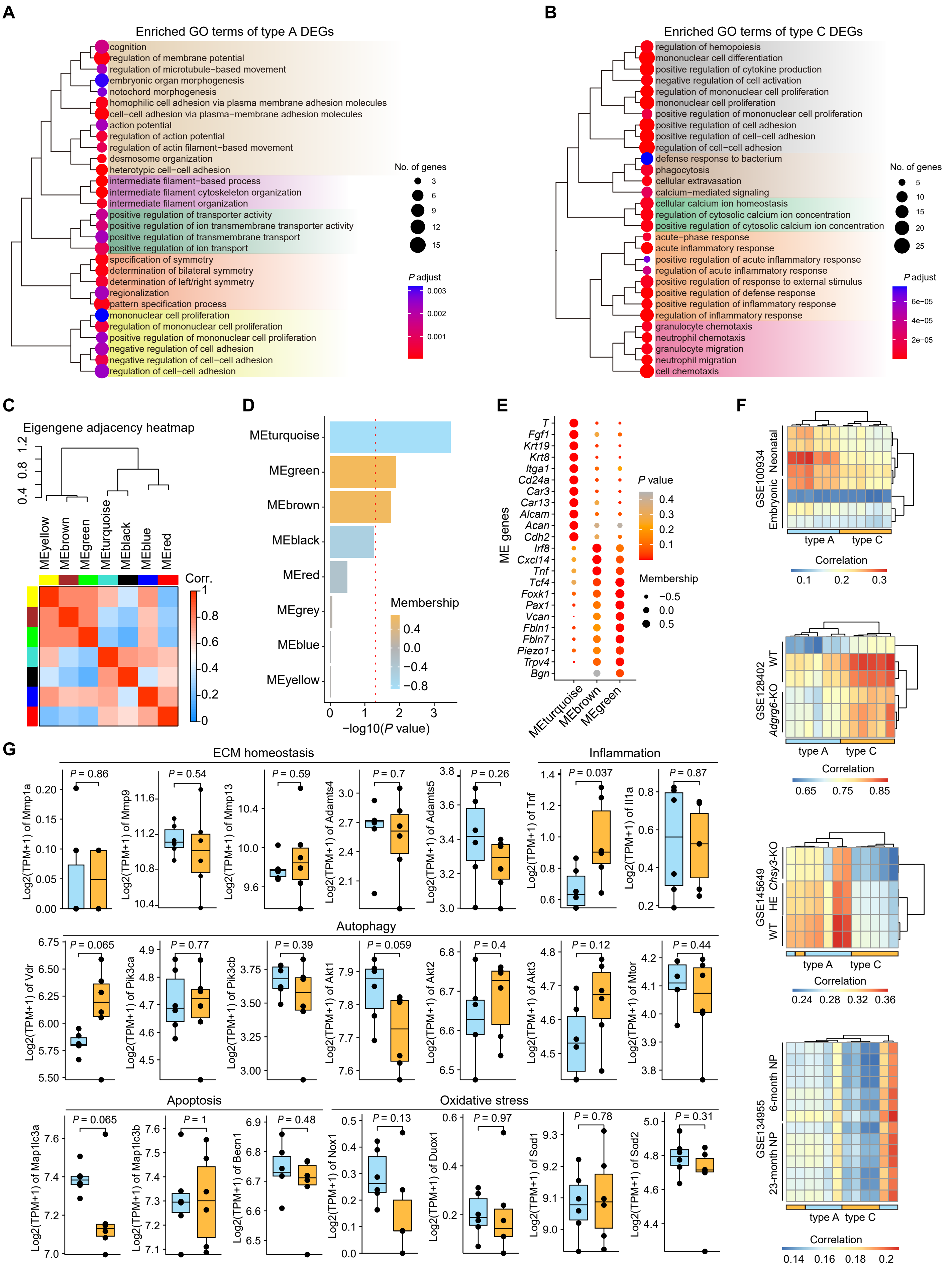

A
Enriched GO terms of type A DEGs
cognition
regulation of membrane potential
regulation of microtubule−based movement
embryonic organ morphogenesis
notochord morphogenesis
homophilic cell adhesion via plasma membrane adhesion molecules
cell−cell adhesion via plasma−membrane adhesion molecules
action potential
regulation of action potential
regulation of actin filament−based movement
desmosome organization
heterotypic cell−cell adhesion
intermediate filament−based process
intermediate filament cytoskeleton organization
intermediate filament organization
positive regulation of transporter activity
positive regulation of ion transmembrane transporter activity
positive regulation of transmembrane transport
positive regulation of ion transport
specification of symmetry
determination of bilateral symmetry
determination of left/right symmetry
regionalization
pattern specification process
mononuclear cell proliferation
regulation of mononuclear cell proliferation
positive regulation of mononuclear cell proliferation
negative regulation of cell adhesion
negative regulation of cell−cell adhesion
regulation of cell−cell adhesion
No. of genes
3
6
9
12
15
P adjust
0.003
0.002
0.001
B
Enriched GO terms of type C DEGs
regulation of hemopoiesis
mononuclear cell differentiation
positive regulation of cytokine production
negative regulation of cell activation
regulation of mononuclear cell proliferation
mononuclear cell proliferation
positive regulation of mononuclear cell proliferation
positive regulation of cell adhesion
positive regulation of cell−cell adhesion
regulation of cell−cell adhesion
defense response to bacterium
phagocytosis
cellular extravasation
calcium−mediated signaling
cellular calcium ion homeostasis
regulation of cytosolic calcium ion concentration
positive regulation of cytosolic calcium ion concentration
acute−phase response
acute inflammatory response
positive regulation of acute inflammatory response
regulation of acute inflammatory response
positive regulation of response to external stimulus
positive regulation of defense response
positive regulation of inflammatory response
regulation of inflammatory response
granulocyte chemotaxis
neutrophil chemotaxis
granulocyte migration
neutrophil migration
cell chemotaxis
No. of genes
5
10
15
20
25
P adjust
6e−05
4e−05
2e−05
C
Eigengene adjacency heatmap
MEyellow
MEbrown
MEgreen
MEturquoise
MEblack
MEblue
MEred
Corr.
D
MEturquoise
MEgreen
MEbrown
MEblack
MEred
MEgrey
MEblue
MEyellow
Membership
−log10(P value)
E
ME genes
T
Fgf1
Krt19
Krt8
Itga1
Cd24a
Car3
Car13
Alcam
Acan
Cdh2
Irf8
Cxcl14
Tnf
Tcf4
Foxk1
Pax1
Vcan
Fbln1
Fbln7
Piezo1
Trpv4
Bgn
MEturquoise
MEbrown
MEgreen
P value
Membership
F
GSE100934
Neonatal
Embryonic
type A
type C
Correlation
GSE128402
WT
Adgrg6-KO
GSE145649
Chsy3-KO
HE
WT
GSE134955
6-month NP
23-month NP
G
ECM homeostasis
Inflammation
Autophagy
Apoptosis
Oxidative stress
Log2(TPM+1) of Mmp1a
Log2(TPM+1) of Mmp9
Log2(TPM+1) of Mmp13
Log2(TPM+1) of Adamts4
Log2(TPM+1) of Adamts5
Log2(TPM+1) of Tnf
Log2(TPM+1) of Il1a
Log2(TPM+1) of Vdr
Log2(TPM+1) of Pik3ca
Log2(TPM+1) of Pik3cb
Log2(TPM+1) of Akt1
Log2(TPM+1) of Akt2
Log2(TPM+1) of Akt3
Log2(TPM+1) of Mtor
Log2(TPM+1) of Map1lc3a
Log2(TPM+1) of Map1lc3b
Log2(TPM+1) of Becn1
Log2(TPM+1) of Nox1
Log2(TPM+1) of Duox1
Log2(TPM+1) of Sod1
Log2(TPM+1) of Sod2
P = 0.86
P = 0.54
P = 0.59
P = 0.7
P = 0.26
P = 0.037
P = 0.87
P = 0.065
P = 0.77
P = 0.39
P = 0.059
P = 0.4
P = 0.12
P = 0.44
P = 0.065
P = 1
P = 0.48
P = 0.13
P = 0.97
P = 0.78
P = 0.31

**Supplementary Figure 3. RNA-seq analysis of type A and type C discs.**

(**A**) and (**B**). Enriched GO term of DEGs in type A (**A**) and type C (**B**) discs. Dot size and colormap indicate the number of genes and enriched p values in each term. Terms are hierarchically clustered and shown in colored shadows.

(**C**) Adjacency heatmap shows correlations among gene modules defined using WGCNA analysis of genes associated with types of discs.

(**D**) Bar chart of module membership (MM), a measure of intra-modular connectivity of each module. Dashed line indicates the *P* value of 0.05.

(**E**) Dot plots show the *P* values (color) and membership (dot size) of ME genes in the three significant modules.

(**F**) Hierarchically-clustered heatmaps depict the correlation between RNA-seq datasets obtained in this study and publicly available datasets. Pearson correlations are computed based on overall gene expressions. Type A discs demonstrate stronger correlations with neonatal discs, while Type C discs do not show any correlations with degenerative discs induced by *Adgrg6*-KO and *Chsy3*-KO, nor do they exhibit correlations with discs with aged NP.

(**G**) Boxplots depict the expression levels of validated marker genes associated with degeneration in two disc types, encompassing ECM hemostasis, inflammation, autophagy, apoptosis, and oxidative stress. With the exception of the inflammatory gene Tnf, no upregulation of these genes is evident in type C discs.

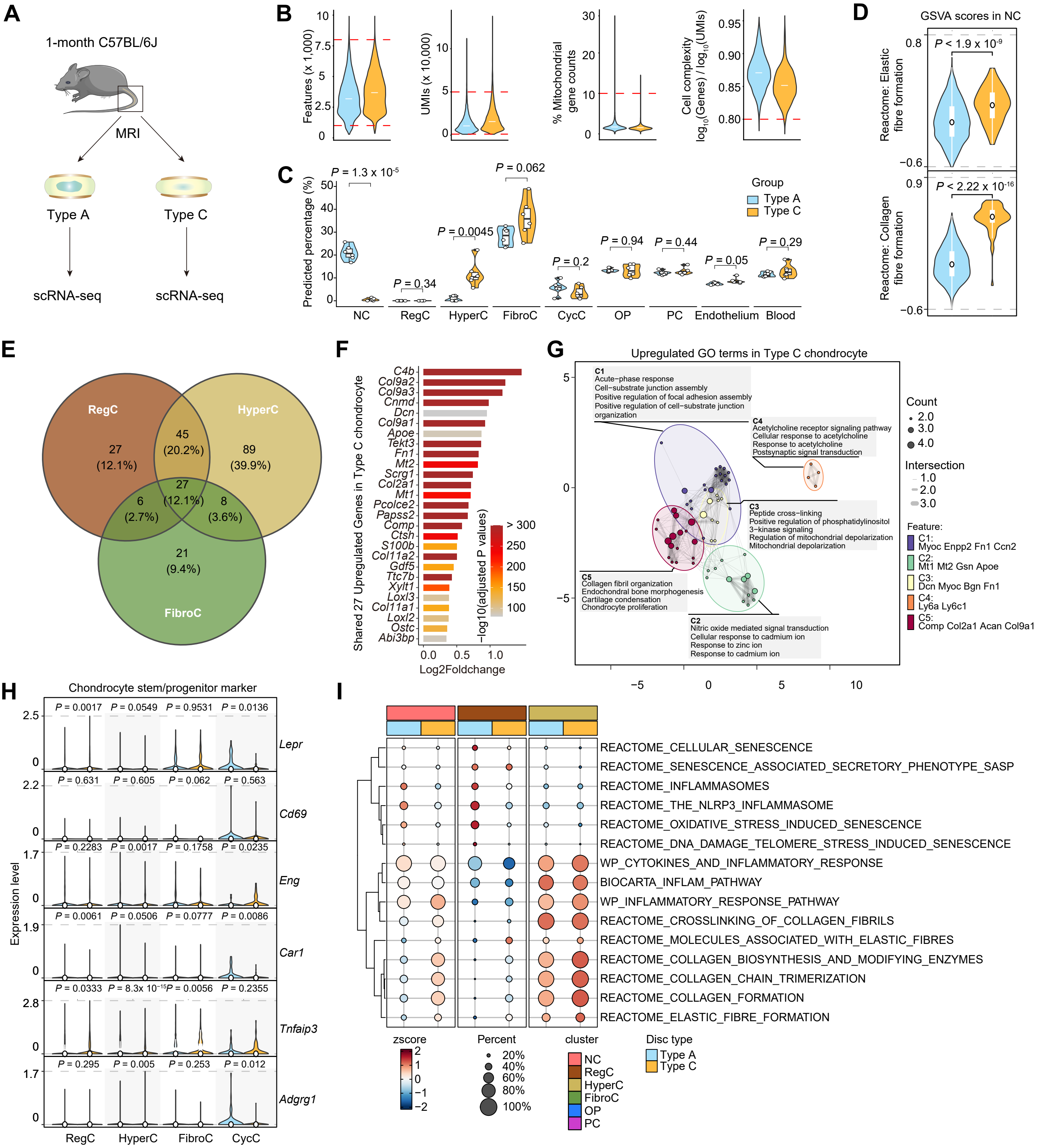
A
1-month C57BL/6J
MRI
Type A
Type C
scRNA-seq
scRNA-seq
B
Features (x 1,000)
UMIs (x 10,000)
% Mitochondrial gene counts
Cell complexity log10(Genes) / log10(UMIs)
C
Predicted percentage (%)
P = 1.3 x 10-5
P = 0.062
P = 0.0045
P = 0.34
P = 0.2
P = 0.94
P = 0.44
P = 0.05
P = 0.29
Group
Type A
Type C
NC
RegC
HyperC
FibroC
CycC
OP
PC
Endothelium
Blood
D
GSVA scores in NC
P < 1.9 x 10-9
Reactome: Elastic fibre formation
P < 2.22 x 10-16
Reactome: Collagen fibre formation
E
RegC
HyperC
FibroC
27 (12.1%)
45 (20.2%)
89 (39.9%)
27 (12.1%)
6 (2.7%)
8 (3.6%)
21 (9.4%)
F
Shared 27 Upregulated Genes in Type C chondrocyte
C4b
Col9a2
Col9a3
Cnmd
Dcn
Col9a1
Apoe
Tekt3
Fn1
Mt2
Scrg1
Col2a1
Mt1
Pcolce2
Papss2
Comp
Ctsh
S100b
Col11a2
Gdf5
Ttc7b
Xylt1
Loxl3
Col11a1
Loxl2
Ostc
Abi3bp
−log10(adjusted P values)
> 300
250
200
150
100
Log2Foldchange
G
Upregulated GO terms in Type C chondrocyte
C1
Acute−phase response
Cell−substrate junction assembly
Positive regulation of focal adhesion assembly
Positive regulation of cell−substrate junction organization
C4
Acetylcholine receptor signaling pathway
Cellular response to acetylcholine
Response to acetylcholine
Postsynaptic signal transduction
C3
Peptide cross−linking
Positive regulation of phosphatidylinositol 3−kinase signaling
Regulation of mitochondrial depolarization
Mitochondrial depolarization
C5
Collagen fibril organization
Endochondral bone morphogenesis
Cartilage condensation
Chondrocyte proliferation
C2
Nitric oxide mediated signal transduction
Cellular response to cadmium ion
Response to zinc ion
Response to cadmium ion
Count
Intersection
Feature:
C1: Myoc Enpp2 Fn1 Ccn2
C2: Mt1 Mt2 Gsn Apoe
C3: Dcn Myoc Bgn Fn1
C4: Ly6a Ly6c1
C5: Comp Col2a1 Acan Col9a1
H
Chondrocyte stem/progenitor marker
Expression level
Lepr
Cd69
Eng
Car1
Tnfaip3
Adgrg1
RegC
HyperC
FibroC
CycC
I
REACTOME_CELLULAR_SENESCENCE
REACTOME_SENESCENCE_ASSOCIATED_SECRETORY_PHENOTYPE_SASP
REACTOME_INFLAMMASOMES
REACTOME_THE_NLRP3_INFLAMMASOME
REACTOME_OXIDATIVE_STRESS_INDUCED_SENESCENCE
REACTOME_DNA_DAMAGE_TELOMERE_STRESS_INDUCED_SENESCENCE
WP_CYTOKINES_AND_INFLAMMATORY_RESPONSE
BIOCARTA_INFLAM_PATHWAY
WP_INFLAMMATORY_RESPONSE_PATHWAY
REACTOME_CROSSLINKING_OF_COLLAGEN_FIBRILS
REACTOME_MOLECULES_ASSOCIATED_WITH_ELASTIC_FIBRES
REACTOME_COLLAGEN_BIOSYNTHESIS_AND_MODIFYING_ENZYMES
REACTOME_COLLAGEN_CHAIN_TRIMERIZATION
REACTOME_COLLAGEN_FORMATION
REACTOME_ELASTIC_FIBRE_FORMATION
zscore
Percent
20%
40%
60%
80%
100%
cluster
NC
RegC
HyperC
FibroC
OP
PC
Disc type
Type A
Type C

**Supplementary Figure 4. scRNA-seq analysis of type A and type C discs.**

(**A**) Schematic of scRNA-seq study design. Mice underwent MRI examinations to determine the classification of discs ranging from Co2/3 to Co7/8. Subsequently, type A and type C discs were extracted, pooled together, and processed to generate cell suspensions for scRNA-seq analysis, respectively.

(**B**) Criterion for quality control of the scRNA-seq dataset is denoted by the dashed lines.

(**C**) Violin plots display the cell percentages of clusters identified in scRNA-seq datasets within each bulk RNA-seq sample, as predicted through Bayesian integrative analysis. P values of significance are determined by Student's t-test.

(**D**) Violin plots show GSVA scores of geneset representing inflammation and fiber synthesis in two types of discs. P values are determined using two tailed Student's t-test.

(**E**) Venn diagram illustrates the quantities and proportions (in parentheses) of upregulated DEGs within three primarily
chondrocyte clusters in type C intervertebral discs.

(**F**) Barplots display 27 shared upregulated DEGs.

(**G**) Enrichmap displays five categories of enriched terms derived from DEGs in type C chondrocytes. The size of each dot corresponds to the number of occurrences of individual genes within the enriched terms. The thickness of the lines represents the strength of intersection between genes within the same category. Different colors on the map distinguish feature genes belonging to distinct categories.

(**H**) Expression of AF differentiation maker genes in chondrocyte clusters. Significance between type A and type C were determined. *P* values were calculated using two-tailed Student's t-tests.

(**I**) Hierarchically-clustered heatmaps depict the gene expression levels associated with inflammation and elastic fibrosis in cell
clusters isolated from type A and type C discs.

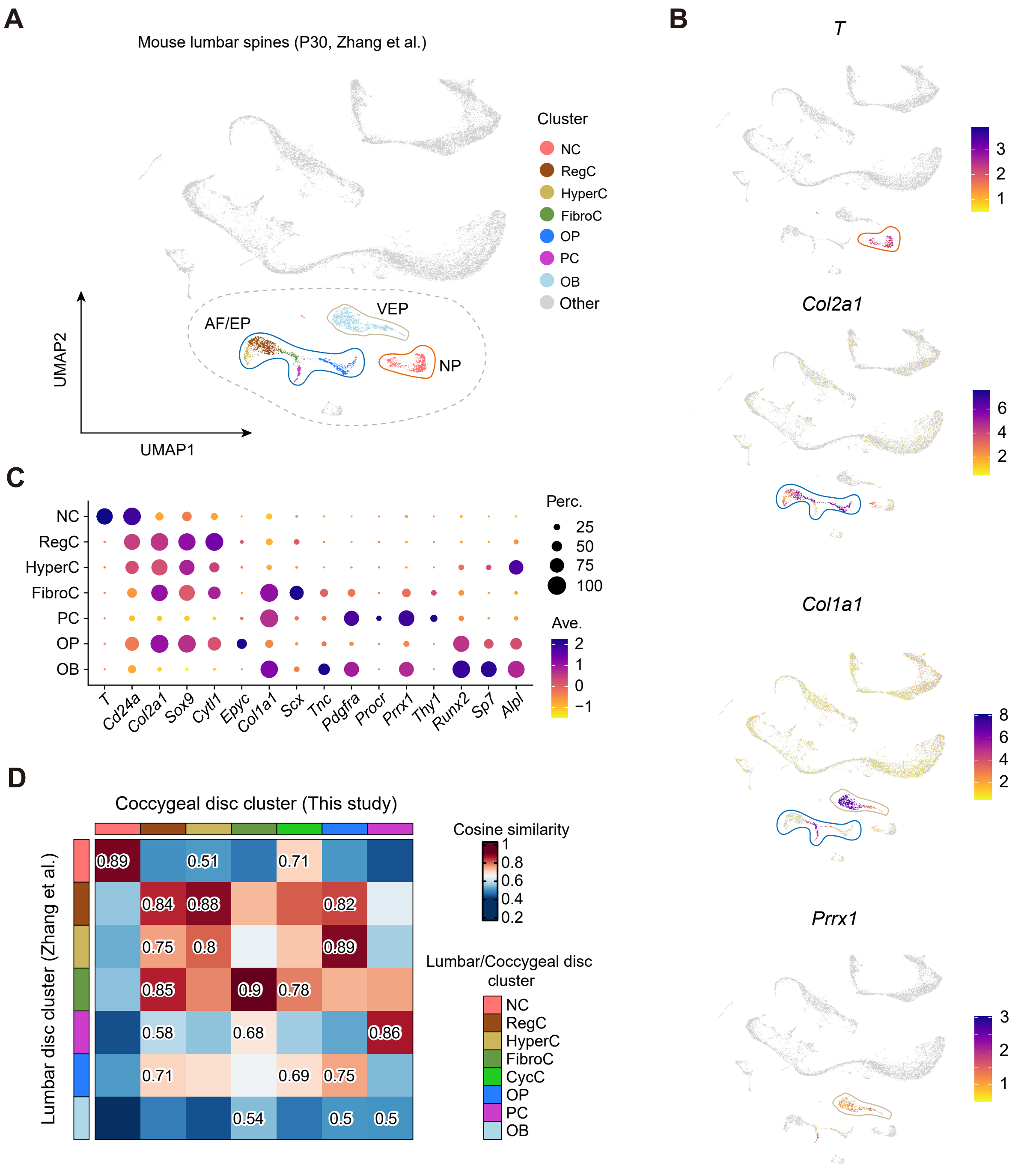

A
Mouse lumbar spines (P30, Zhang et al.)
Cluster
NC
RegC
HyperC
FibroC
OP
PC
OB
Other
VEP
AF/EP
NP
UMAP2
UMAP1
B
T
Col2a1
Col1a1
Prrx1
C
NC
RegC
HyperC
FibroC
PC
OP
OB
T
Cd24a
Col2a1
Sox9
Cytl1
Epyc
Col1a1
Scx
Tnc
Pdgfra
Procr
Prrx1
Thy1
Runx2
Sp7
Alpl
Perc.
25
50
75
100
Ave.
2
1
0
−1
D
Coccygeal disc cluster (This study)
Lumbar disc cluster (Zhang et al.)
Cosine similarity
1
0.8
0.6
0.4
0.2
Lumbar/Coccygeal disc cluster
NC
RegC
HyperC
FibroC
CycC
OP
PC
OB
0.89
0.51
0.71
0.84
0.88
0.82
0.75
0.8
0.89
0.85
0.9
0.78
0.58
0.68
0.86
0.71
0.69
0.75
0.54
0.5
0.5

**Supplementary Figure 5. Re-analysis of scRNA-seq data of mouse lumbar spine cells.**

(**A**) Re-analysis of scRNA-seq dataset from mouse lumbar spine at P30 (GSE235198).

(**B**) NP, AF/EP, and VB cell clusters are in-silico inferred by the expression of *T* (marker for Notochord cell), *Col2a1* (marker for chondrocyte), *Col1a1* (marker for fibroblast and osteogenic cell), and *Prrx1* (marker for skeletal cells in vertebra and bone).

(**C**) Expression of curated genes for cell cluster annotation.

(**D**) Heatmaps display the pairwise similarities between mouse lumbar disc cell clusters and type A and type C disc cell clusters. Correlations are calculated using highly variable genes in each dataset, with the distance metric set to "cosine".

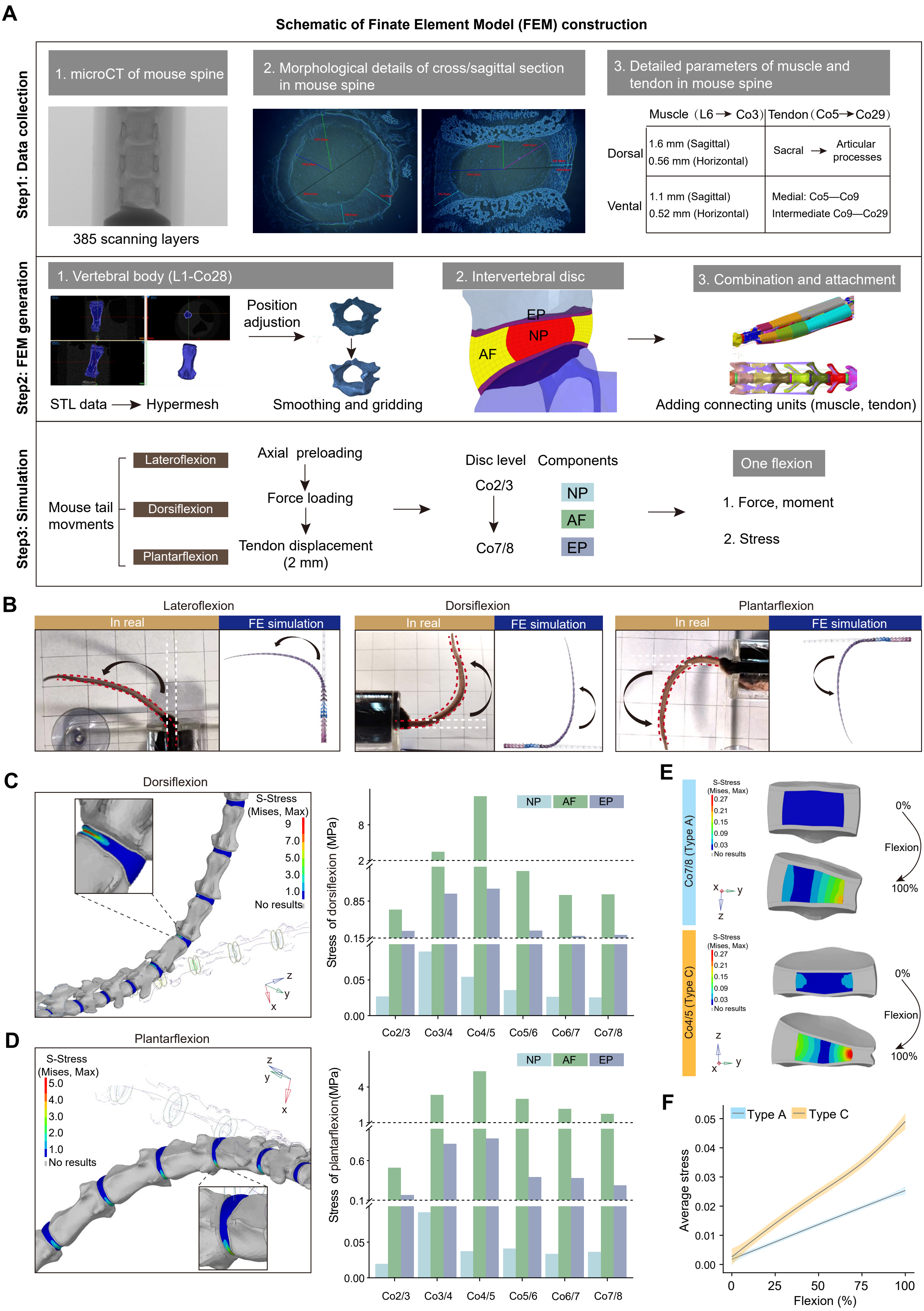
A
Schematic of Finate Element Model (FEM) construction
Step1: Data collection
1. microCT of mouse spine
385 scanning layers
2. Morphological details of cross/sagittal section in mouse spine
3. Detailed parameters of muscle and tendon in mouse spine
Muscle (L6 → Co3)
Tendon (Co5 → Co29)
Dorsal
1.6 mm (Sagittal)
0.56 mm (Horizontal)
Sacral → Articular processes
Vental
1.1 mm (Sagittal)
0.52 mm (Horizontal)
Medial: Co5—Co9
Intermediate Co9—Co29
Step2: FEM generation
1. Vertebral body (L1-Co28)
STL data → Hypermesh
Position adjustion
Smoothing and gridding
2. Intervertebral disc
EP
NP
AF
3. Combination and attachment
Adding connecting units (muscle, tendon)
Step3: Simulation
Mouse tail movments
Lateroflexion
Dorsiflexion
Plantarflexion
Axial preloading
Force loading
Tendon displacement (2 mm)
Disc level
Co2/3
Co7/8
Components
NP
AF
EP
One flexion
1. Force, moment
2. Stress
B
Lateroflexion
Dorsiflexion
Plantarflexion
In real
FE simulation
C
Dorsiflexion
S-Stress (Mises, Max)
9
7.0
5.0
3.0
1.0
No results
Stress of dorsiflexion (MPa)
NP AF EP
Co2/3 Co3/4 Co4/5 Co5/6 Co6/7 Co7/8
D
Plantarflexion
S-Stress (Mises, Max)
5.0
4.0
3.0
2.0
1.0
No results
Stress of plantarflexion(MPa)
E
Co7/8 (Type A)
Co4/5 (Type C)
S-Stress (Mises, Max)
0.27
0.21
0.15
0.09
0.03
No results
0%
Flexion
100%
F
Type A
Type C
Average stress
Flexion (%)

**Supplementary Figure 6. Construction strategy and flexion simulations of finite element model (FEM) of mouse spine**

(**A**) Generalizable framework of FEM construction of mouse tail spine. Three steps including biological data collection, FEM computational generation, and flexion simulation are described in detail.

(**B**) Photographs illustrate the lateroflexion (left), dorsiflexion (middle), and plantarflexion (right) of the mouse tail. The curved arrow indicates the direction of flexion. The white dashed line indicates the start/relaxing position and the red dashed line indicates the end/flexion position, along with simulations of the three flexion processes using mouse spine FEM.

(**C**) **and** (**D**) View of von Mises stress distributed in AF of mouse tail discs during dorsiflexion (**C**) and plantarflexion (**D**). Magnified view showing the stress distribution of Co4/5 disc. Bar charts show the stress of NP, AF, and EP at the end of lateroflexion.

(**E**) View of von Mises stress distributed in NP between type A and type C discs at the start and end of flexion.

(**F**) Smooth curves indicate linear regression 95% confidence interval of the average stress in NP of two types of discs (type A: Co2/3, Co6/7, and Co7/8; type C: Co3/4, Co4/5, and Co5/6).

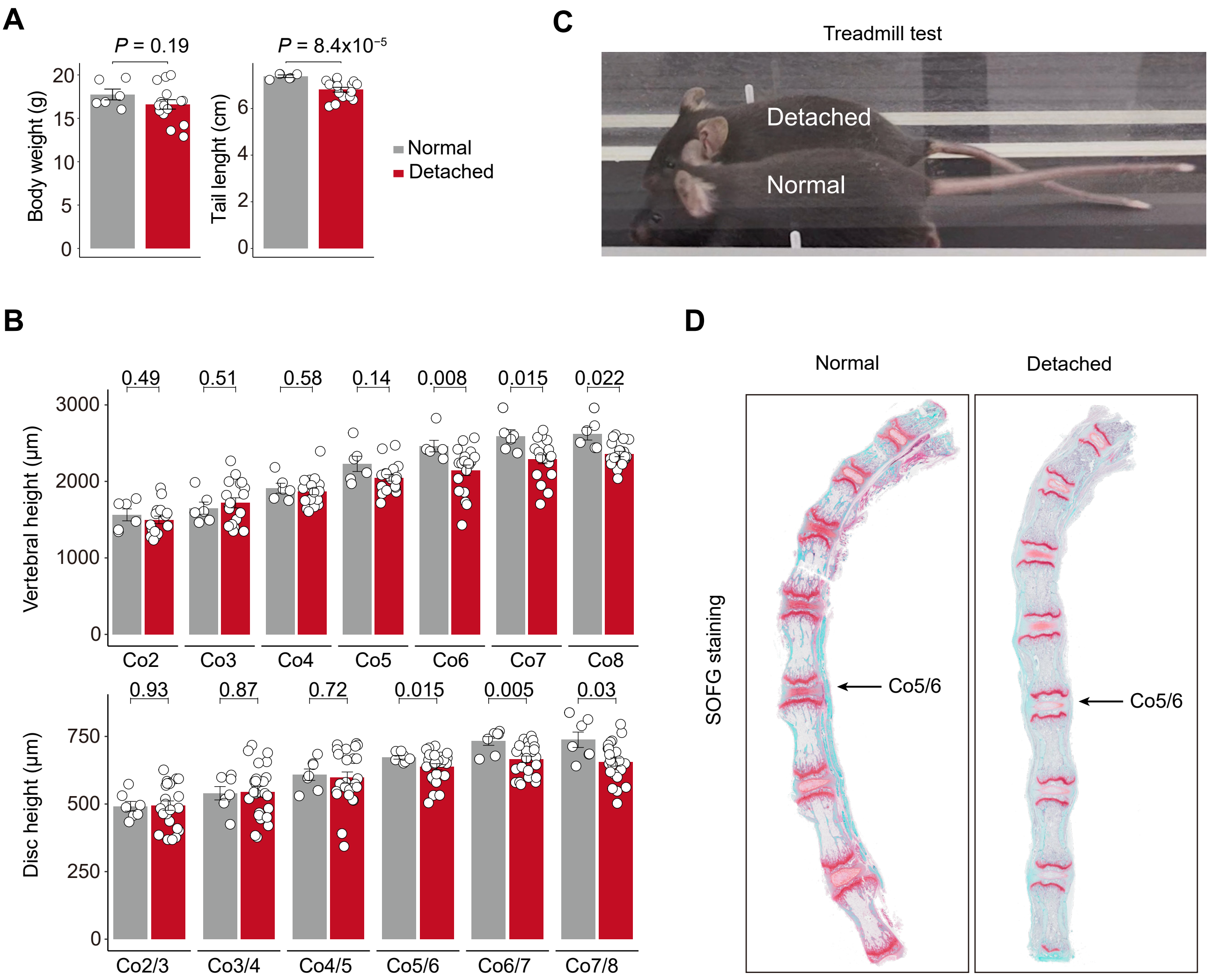
A
P = 0.19
P = 8.4x10−5
Body weight (g)
Tail lenght (cm)
Normal
Detached
C
Treadmill test
Detached
Normal
B
0.49
0.51
0.58
0.14
0.008
0.015
0.022
Vertebral height (μm)
Co2
Co3
Co4
Co5
Co6
Co7
Co8
0.93
0.87
0.72
0.015
0.005
0.03
Disc height (μm)
Co2/3
Co3/4
Co4/5
Co5/6
Co6/7
Co7/8
D
Normal
Detached
SOFG staining
Co5/6
Co5/6

**Supplementary Figure 7. Tendon detachment prevent the formation of the type C discs.**

(**A**) Bar chart show the body weight (left) and tail length (right) of mice from the normal control with sham operation (N = 6) and tendon detachment group (N = 15). Data are mean ± S.E. *P* values are determined using two tailed Student's t-test.

(**B**) Representative photograph depicts a treadmill test reveals that mice in the normal control group exhibit well-coordinated tail wagging, whereas those with tendon detachment display a lack of control over their tail movements.

(**C**) Bar chart show the height of vertebrae ranging from Co2 to Co8 (top) and height of disc ranging from Co2/3 to Co7/8 (bottom) between normal (N = 7) and detached groups (N = 24). Data are mean ± S.E. *P* values are determined using two tailed Student's t-test.

(**D**) Representative SOFG staining on the mouse spine ranging from Co1 to Co8, revealed distinct features in the Co5/6 discs of the normal and detached groups. Specifically, the Co5/6 disc in the normal group displays type C characteristics, whereas the Co5/6 disc in the detached group exhibits type A features.

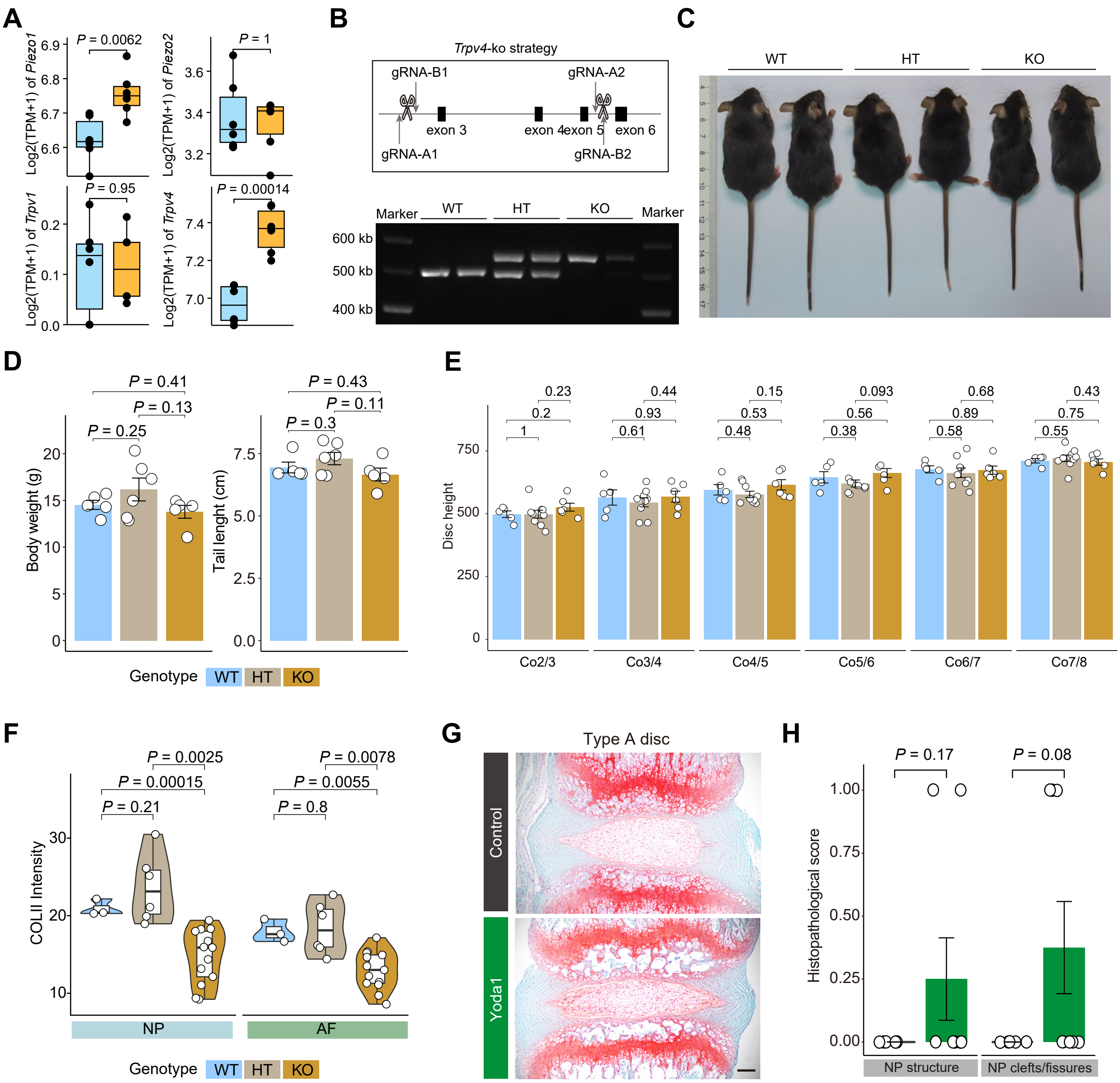

A
B
C
D
E
F
G
H
Trpv4-ko strategy
gRNA-B1
gRNA-A2
exon 3
exon 4
exon 5
exon 6
gRNA-A1
gRNA-B2
Marker
WT
HT
KO
600 kb
500 kb
400 kb
Body weight (g)
Tail lenght (cm)
Disc height
Co2/3
Co3/4
Co4/5
Co5/6
Co6/7
Co7/8
Genotype
COLII Intensity
NP
AF
Type A disc
Control
Yoda1
Histopathological score
NP structure
NP clefts/fissures

**Supplementary Figure 8. Trpv4-knockout reduce the type C characteristics**

(**A**) Boxplots depict the expression levels of genes associated with mechanosensation in two disc types. *P* values of significance are determined by Wilcoxon rank-sum test.

(**B**) Construction strategy for Trpv4-KO mice using utilizing CRISPR-Cas9 technology by deleting three exons guided by two pairs of gRNAs (top). The efficiency of the knockout is verified through PCR validation (bottom).

(**C**) Dorsal views of wildtype (WT), heterozygote (HT), and knockout (KO) mice.

(**D**) Bar chart show the body weight (left) and tail length (right) of mice from the WT (N = 5), HT (N = 6), and KO (N = 5). Data are mean ± S.E. *P* values are determined using Wilcoxon rank-sum test.

(**E**) Bar chart show the height of height of disc ranging from Co2/3 to Co7/8 among WT (N = 5), HT (N = 9), and KO (N = 9). Data are mean ± S.E. *P* values are determined using Wilcoxon rank-sum test.

(**F**) Quantification of signaling density of COL II in NP region (left) and AF region (right) among WT (N = 3), HT (N = 6), and KO (N = 13). Scale bar, 100 μm. *P* values are determined using Wilcoxon rank-sum test.

(**G**) Representative SOFG staining show the morphology changes of NP in type A disc of mice injected with Yoda1. Scale bar, 100 μm. Histological scores of type A discs from control and injection group. Data are mean ± S.E. *P* values are determined by two tailed Student's t-tests.
